# Reference Repulsion Is Not a Perceptual Illusion

**DOI:** 10.1101/366393

**Authors:** Matthias Fritsche, Floris P. de Langea

## Abstract

Perceptual decisions are often influenced by contextual factors. For instance, when engaged in a visual discrimination task against a reference boundary, subjective reports about the judged stimulus feature are biased away from the boundary – a phenomenon termed reference repulsion. Until recently, this phenomenon has been thought to reflect a perceptual illusion regarding the appearance of the stimulus, but new evidence suggests that it may rather reflect a post-perceptual decision bias. To shed light on this issue, we examined whether and how orientation judgments affect perceptual appearance. In a first experiment, we confirmed that after judging a grating stimulus against a discrimination boundary, the subsequent reproduction response was indeed repelled from the boundary. To investigate the perceptual nature of this bias, in a second experiment we measured the perceived orientation of the grating stimulus more directly, in comparison to a reference stimulus visible at the same time. Although we did observe a small repulsive bias away from the boundary, this bias was explained by random trial-by-trial fluctuations in sensory representations together with classical stimulus adaptation effects and did not reflect a systematic bias due to the discrimination judgment. Overall, the current study indicates that discrimination judgments do not elicit a perceptual illusion and points towards a post-perceptual locus of reference repulsion.

## 1. Introduction

Perceptual decisions are influenced by a multitude of contextual factors such as recent sensory input (Snyder, Schwiedrzik, Vitela, & Melloni, 2015), previous decisions (Akaishi, Umeda, Nagase, & Sakai, 2014), rewards (Mulder, Wagenmakers, Ratcliff, Boekel, & Forstmann, 2012) and prior expectations (Summerfield & de Lange, 2014). Importantly, some of these contextual biases measured in participants’ responses arise early during visual processing, for instance during the encoding of the physical stimulus or decoding of the sensory representation (Webster, 2015), whereas others arise at later post-perceptual decisional stages (Firestone & Scholl, 2016; Fritsche, Mostert, & de Lange, 2017), during working memory retention (Bliss, Sun & D’Esposito, 2017; Huang & Sekuler, 2010; Papadimitriou, Ferdoash & Snyder, 2015; Visscher, Kahana, & Sekuler, 2009) or when a motor response is formed (Pape & Siegel, 2016). Consequently, some contextual biases affect the perceptual appearance of a sensory stimulus, eliciting perceptual illusions, while others only affect the decision or response after the stimulus has been perceived. The stage at which a bias arises has important implications, as it determines what information the organism has available at any given point in time. For instance, a bias that is introduced early during visual processing, affecting perceptual appearance, would be carried through all representations on subsequent processing stages and would lead to persistent behavioral biases, regardless at which point in time the representation is read out for behavior and in which particular way the representation is probed. In contrast, a bias that only arises at later stages, for example during working memory retention, could be modulated by deferred factors such as working memory interference or the duration of the retention interval before a behavior is produced (e.g. Bliss et al., 2017). Consequently, perceptual and post-perceptual biases can lead to dramatically different behavioral outcomes, depending on the situation in which a perceptual decision is probed. In order to develop a full understanding of human perceptual decision making it is therefore vital to understand at which processing stage a bias arises. Furthermore, insights into the stage at which a bias arises will inform the search about the neural substrates and mechanisms underlying these phenomena.

A bias that has been previously argued to affect the appearance of a visual stimulus is so-called reference repulsion. Reference repulsion refers to the phenomenon that subjective estimates about a stimulus attribute are biased away from an external reference, against which the stimulus attribute has been compared. For instance, in a famous study by Jazayeri & Movshon (2007) participants had to judge whether the motion direction of a random-dot motion stimulus was clockwise or counterclockwise of a reference line that formed a discrimination boundary. On a subset of trials participants subsequently had to reproduce the average motion direction. Jazayeri and Movshon found that reproduction responses were systematically biased away from the discrimination boundary, and that this bias was stronger for noisier stimuli. This bias was interpreted as stemming from a sensory decoding strategy with a bimodal weighting profile that is optimized for the clockwise/counterclockwise discrimination task. Crucially, this theory assumes that reference repulsion occurs as a consequence of being engaged in the discrimination judgment task and arises *during* perceptual processing, therefore resulting in a perceptual illusion. An alternative theory proposed that the repulsion could arise from Bayesian inference with the initial discrimination judgment constraining the prior over motion directions when forming a perceptual decision about the overall motion direction (Stocker & Simoncelli, 2008; Luu & Stocker, 2018). Under these theories it is not clear whether reference repulsion would occur at a perceptual (Stocker & Simoncelli, 2008) or post-perceptual stage (Luu & Stocker, 2018).

A recent study cast doubt on the perceptual nature of reference repulsion (Zamboni, Ledgeway, McGraw, & Schluppeck, 2016). Although replicating the original results, Zamboni et al. found that the presence of the discrimination boundary during the reproduction phase was necessary for reference repulsion to occur. When the discrimination boundary was presented only during the time of stimulus presentation, when participants made the fine discrimination judgment, subsequent stimulus reproductions appeared to be unbiased, suggesting that participants could access a veridical representation of the stimulus. The authors proposed that the discrimination boundary presented during the reproduction phase served as an anchor that, when combined with sensory evidence, led to a late, decision-related bias.

Does reference repulsion therefore reflect a purely post-perceptual, decision-related phenomenon, while the appearance of the sensory stimulus is unbiased? Although Zamboni et al.’s study suggests that the large biases previously seen in stimulus reproductions are decision-related, their analysis might not have been sensitive enough to reveal smaller perceptual reference repulsion biases. This is corroborated by a recent re-analysis of Zamboni et al.’s data (Luu & Stocker, 2018), which revealed that the data were significantly better fit by a model incorporating reference repulsion than by a model without such a bias. The re-analysis by Luu & Stocker suggests that reproduction responses are biased, albeit more weakly, even when the discrimination boundary is absent during the reproduction phase. While this revives the possibility that reference repulsion may affect the appearance of visual stimuli, reproduction responses may heavily factor in post-perceptual processes and therefore it is not clear whether reference repulsion is a perceptual illusion. In order to shed light on this issue, we first examined whether reference repulsion in reproduction responses would occur even when the discrimination boundary was presented only before the presentation of the actual stimulus and crucially not during the reproduction phase, as suggested by Luu & Stocker’s reanalysis of the experiment by Zamboni et al. Focusing on orientation perception, for which similar biases as in motion perception have been reported (Luu & Stocker, 2018), we indeed found a robust reference repulsion bias. Next, we investigated the perceptual nature of this bias by using a more direct comparison technique to measure perception. In particular, we measured the perceived orientation of a probe stimulus, in direct comparison to a reference stimulus visible at the same time, after the probe stimulus was judged against a discrimination boundary. In contrast to the reproduction technique, in which stimuli are reproduced on the basis of a memory representation, rendering responses susceptible to post-perceptual working memory and decision biases, the technique of a direct perceptual comparison between simultaneously presented probe and reference stimuli reduces the influence of such post-perceptual processes and has the sensitivity to reveal subtle perceptual biases (Fritsche et al., 2017). To foreshadow, we found a repulsive bias away from the boundary. However, this bias was markedly smaller than the bias in reproduction responses and, importantly, could be completely explained by random trial-by-trial fluctuations in sensory representations together with classical negative stimulus adaptation and thus did not reflect a systematic bias due to the discrimination judgment. We conclude that rather than eliciting a perceptual illusion, making discrimination judgments leads to a post-perceptual bias in decisions or working memory representations.

## 2. Experiment 1: Reference repulsion in reproduction responses?

A previous study suggested that reproduction responses are repelled from a discrimination boundary only if the boundary continued to be visible during the reproduction phase of a trial, pointing to a post-perceptual locus of the bias (Zamboni et al., 2016). However, the data analysis might not have been sensitive enough to detect potentially subtler reference repulsion biases when the discrimination boundary was absent during stimulus reproduction (see Luu & Stocker, 2018). Therefore, in Experiment 1 we sought to revisit this finding and test whether reference repulsion could also occur when the discrimination boundary was presented only before the presentation of the actual stimulus, but not during the reproduction phase. Such a bias, which is not dependent on the presence of the boundary during the post-perceptual reproduction phase, might reflect a bias in perceptual appearance.

### 2.1. Methods

#### 2.1.1. Participants

Twenty-four naïve participants (15 female/9 male, age range 19 – 30 years) took part in Experiment 1. All participants reported normal or corrected-to-normal vision and gave written, informed consent prior to the start of the study. The study was approved by the local ethical review board (CMO region Arnhem-Nijmegen, The Netherlands) and was in accordance with the Declaration of Helsinki.

#### 2.1.2. Apparatus and stimuli

Visual stimuli were generated with the Psychophysics Toolbox (Brainard, 1997) for MATLAB (The MathWorks, Natick, MA) and were displayed on a 24” flat panel display (Benq XL2420T, resolution 1920 x 1080, refresh rate: 60Hz). Participants viewed the stimuli from a distance of 53 cm in a dimly lit room, resting their head on a table-mounted chinrest.

A central white fixation dot of 0.25° visual angle diameter was presented on a mid-grey background throughout the whole experiment block. Participants were instructed to maintain fixation at all times.

A prior discrimination boundary was formed by two opposing white line segments (length 0.5°, width 0.09°), presented around the center (10° left or right of fixation) of the upcoming stimulus location, offset by 6.5° (**Figure 1A**).

**Figure 1:**
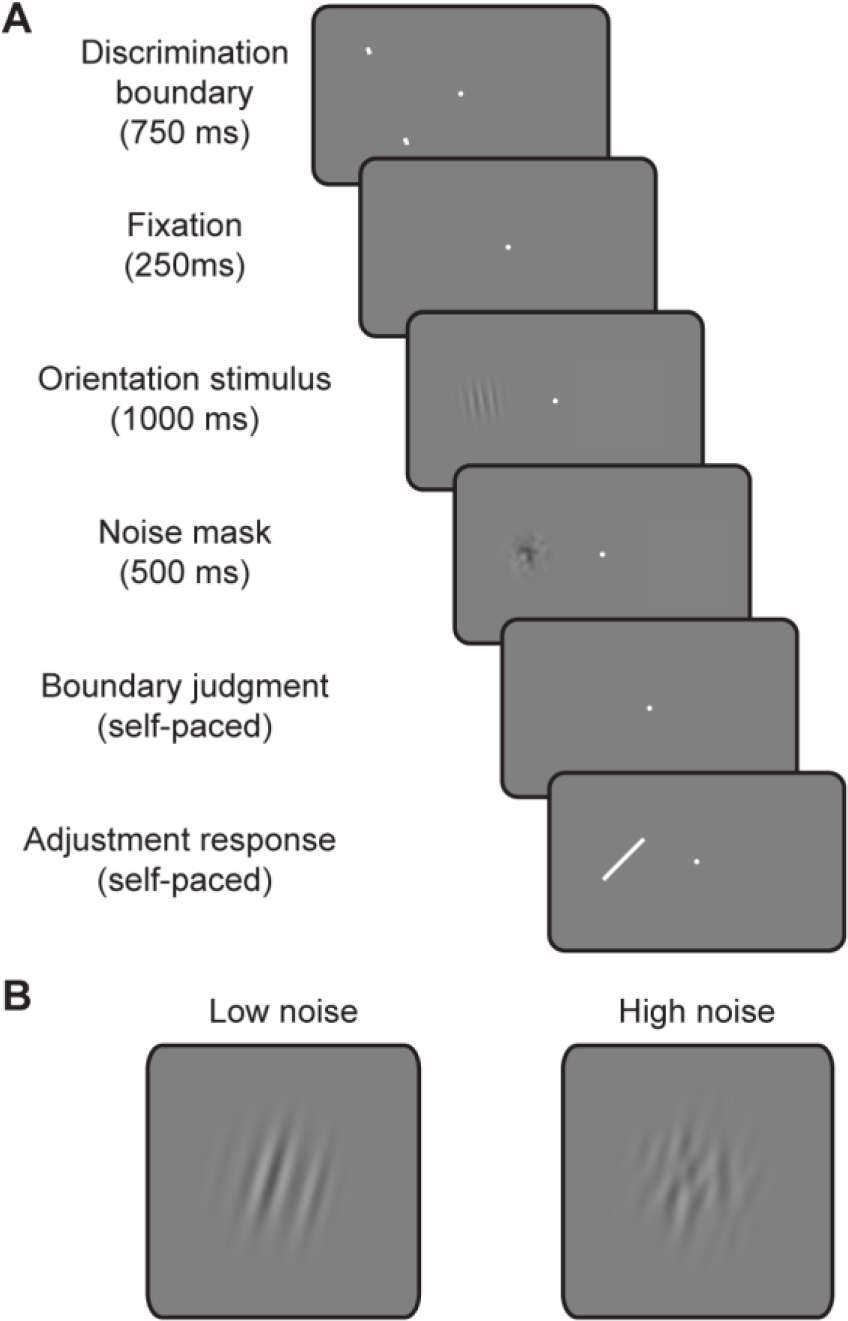
Task and stimuli of Experiment 1. (A) Participants had to indicate whether a grating stimulus was oriented clockwise or counterclockwise with respect to a prior discrimination boundary presented at the same location. Subsequently, participants had to adjust a response bar in order to reproduce the orientation of the grating stimulus. (B) Two predefined noise levels for the grating stimuli were used throughout the experiment. The stimulus on a particular trial either contained low (left, *κ* = 500) or high noise (right, *κ* = 5).

Orientation stimuli were generated by filtering white noise in the Fourier domain with a band-pass filter. The passband of spatial frequencies was defined as a Gaussian with a mean of 0.8 cycles/° and standard deviation of 0.3 cycles/°. The passband for orientations was defined as a von Mises distribution with location parameter *µ* and concentration parameter *κ*. The location parameter *µ* determined the mean orientation of a stimulus, while the concentration parameter *κ* effectively determined the amount of orientation noise. For a high concentration parameter, only few orientations other than the specified mean orientation were present in the signal, resulting in a low noise stimulus. Conversely, a low concentration parameter resulted in a noisy stimulus with a more uniform orientation distribution. Two predetermined noise levels were used in the experiment (**Figure 1B**; low noise: *κ* = 500; high noise: *κ* = 5). After applying the inverse Fourier transform, the root mean square contrast of the stimuli was set to 15.62% of their mean luminance. The stimuli were windowed with a Gaussian envelope (*SD* = 1.5°) and presented at 10° horizontal eccentricity from fixation.

Noise masks were generated by smoothing white noise with a Gaussian kernel (*SD* = 0.3°), windowed by a Gaussian envelope (*SD* = 1.5°) and presented at 50% Michelson contrast at the same location as the orientation stimulus.

Shortly after the offset of the noise mask, a white response bar (length 4°, width 0.09°) was presented at the same location as the orientation and mask stimuli.

#### 2.1.3. Procedure

At the beginning of each trial an oriented discrimination boundary was presented left or right of fixation for 750 ms. The orientation of the discrimination boundary was randomly drawn from the range of all possible orientations, [0-180°). After further 250 ms of fixation, a low or high noise orientation stimulus was presented at the same side as the prior boundary for 1000 ms. The relative orientation of the stimulus with respect to the discrimination boundary was varied from - 15 to 15° in 1° steps. Subsequently, the orientation stimulus was replaced by a noise mask, presented for 500 ms. The noise mask was presented in order to eliminate potential negative afterimages of the orientation stimulus. After the offset of the noise mask participants indicated whether the orientation stimulus was oriented more clockwise or counterclockwise than the prior boundary by pressing the left/right arrow key (boundary judgment). The self-paced response was followed by a 250 ms interval of fixation, after which a white response bar appeared at the same location as the stimulus and mask. Participants were asked to reproduce the orientation of the stimulus by adjusting the response bar with the left and right arrow key (reproduction response). The response was submitted by pressing the space bar. The response was followed by a 1-second inter-trial-interval, before the next trial began.

Participants completed a total of 1,488 trials in two sessions, each split into 8 blocks. Sessions were conducted on different days, but no more than 5 days apart. The trial sequence was counterbalanced with respect to the horizontal location of stimulus presentation, noise level of the stimuli and the relative stimulus orientation with respect to the discrimination boundary. Trials were presented in pseudorandom order.

At the beginning of the first session, participants practiced the boundary judgment task in isolation in blocks of 32 trials at the easiest stimulus level (stimulus orientation 15° clockwise or counterclockwise from boundary) and received trial-by-trial feedback about the correctness of their response via a color change of the fixation dot (red/green). After a participants were sufficiently trained, they then practiced the main task for at least one block of 32 trials. The practice of the main task was repeated at the beginning of the second session.

#### 2.1.4. Data analysis

Psychometric curves were fit to the boundary judgment data of each individual participant, separately for low and high noise conditions. Fits were obtained with the Palamedes toolbox for analyzing psychophysical data (Prins & Kingdom, 2009). The proportion of “clockwise” responses was expressed as a function of the probe stimulus orientation relative to the discrimination boundary (**Figure 2A**). The data were fit with a psychometric function *Ψ*(*x*; *α, β, λ*) = *λ* + (1 – 2*λ*) *F*(*x*; *α, β*), where *Ψ* describes the proportion of “clockwise” responses and *F* is a cumulative Gaussian function with location parameter *α* and slope parameter *β.* Furthermore, *x* denotes the probe stimulus orientation relative to the discrimination boundary and *λ* accounts for stimulus independent lapses. Parameter estimates of low and high noise conditions were statistically compared using two-sided paired *t*-tests at a significance level of *α* = 0.05. We also tested for general biases in low and high noise conditions, reflected in the location parameter *α,* with two-sided one-sample t-tests at a significance level of *α* = 0.05.

**Figure 2:**
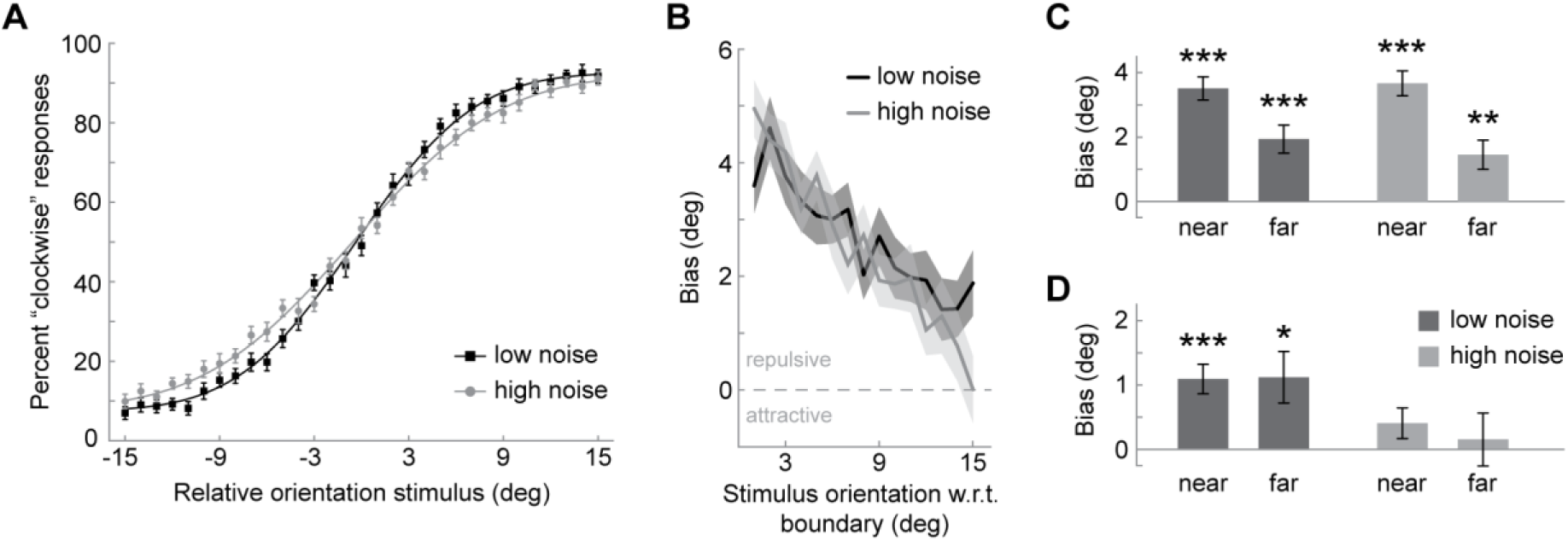
Results of Experiment 1. (A) Boundary judgments: Participants had to indicate whether the grating stimulus was oriented more clockwise or counterclockwise than a prior discrimination boundary. The proportion of “clockwise” responses was expressed as a function of the relative orientation of the stimulus. For positive *x* values the stimulus was oriented more clockwise than the discrimination boundary. The data for low and high noise (black and grey data points) were separately fit with cumulative Gaussian functions (black and grey lines). (B) Reproduction responses: Participants had to reproduce the stimulus orientation by adjusting a response bar. We expressed the signed response error as a function of the orientation difference between stimulus and discrimination boundary. Positive values of bias indicate repulsive errors away from the boundary. Black and grey lines denote low and high noise trials, respectively. Only trials with a correct boundary judgment are considered. (C) We binned the response errors in (B) into near (0-7°) and far (8-15°) orientation distances from the discrimination boundary. Dark and light grey bars represent low and high noise trials, respectively. (D) Same as in (C) but both trials with correct and incorrect boundary judgments are considered. *** p < 0.001, ** p < 0.01, * p < 0.05. Data points represent group means and error bars depict *SEMs*.

In order to investigate whether the reproduction responses were systematically biased due to making fine discrimination judgments, we computed the average signed response error, i.e. the mismatch between the reproduced and actual stimulus orientation, for each relative orientation difference between the stimulus and discrimination boundary. We excluded trials in which the stimulus had the same orientation as the boundary and only considered trials with a correct boundary judgment. Further, we merged the signed response errors for clockwise and counterclockwise stimulus orientations after inverting the sign of response errors to counterclockwise stimuli. This was done in order to quantify the directional response error (away/towards the boundary) as a function of the absolute orientation difference between stimulus and discrimination boundary. In order to simplify the statistical analysis, we separately averaged the response errors for orientations close to the boundary (1 to 7°, near orientations) and further away from the boundary (8 to 15°, far orientations). Subsequently, we conducted a repeated measures ANOVA with the orientation difference between the orientation stimulus and boundary (near/far) and the stimulus noise (low/high) as repeated measures factors.

By considering only correct trials in the above analysis, one is, to some degree, discarding trials in which external and/or internal noise fluctuations in the initial stimulus representation favored a boundary judgment that was opposite to the nominally correct judgment. Conversely, one is more likely to retain trials in which random noise fluctuations biased the internal stimulus representation away from the boundary, towards the correct side of the boundary judgment. Since reproduction responses are likely sensitive to such fluctuations in stimulus representations, selectively retaining trials with correct boundary judgments might therefore introduce an apparent bias in the analysis. In order to ensure that the bias observed in the analysis was not entirely introduced by these random trial-by-trial fluctuations, we conducted a second analysis that was similar to the first, but took all trials into account. As this analysis does not factor in the participants’ boundary judgments it precludes a systematic influence of random trial-by-trial fluctuations and therefore indicates genuine biases in reproduction responses. However, for biases that are caused by the boundary judgment the latter analysis is expected to yield lower estimates, especially for stimuli oriented close to the discrimination boundary and high noise stimuli. This is because boundary judgments for these stimuli are less strongly correlated to the actual stimulus orientation and therefore even a strong bias that is caused by the boundary judgment may cancel out when sorting only according to stimulus orientation. The merit of presenting both analyses will further become evident together with the results of Experiment 2.

### 2.2. Results

The participants’ boundary judgments were meaningfully related to the stimuli and boundaries presented during the experiment (**Figure 2A**). Observers had higher sensitivity for low noise stimuli than for high noise stimuli, which was reflected in a steeper slope of the psychometric curves for the low compared to the high noise condition (low noise *β = 0.1665 +- 0.0098 (SEM),* high noise *β = 0.1328 +- 0.0058 (SEM); t*(23) = 3.4520, *p* = 0.0022). There were no significant general biases in boundary judgments (low noise *α = - 0.3376 +- 0.3205 (SEM), t*(23) = -1.0311, *p =* 0.3132; high noise *α =- 0.5964 +- 0.3080 (SEM), t*(23*) =* -1.8957, *p* = 0.0706) and no significant difference across low and high noise conditions (*t*(23) = 1.6504, *p* = 0.1125). Furthermore, lapse rates were not different across conditions (low noise λ = 0.0529 +- 0.01, high noise λ = 0.0654 +- 0.0106; *t*(23) = -1.1398, *p* = 0.2661).

When investigating biases in reproduction responses, we found that reproductions were clearly repelled from the discrimination boundary and this bias decreased with increasing orientation difference between stimulus and boundary (**Figure 2B**). After binning the data into “near” and “far bins” (**Figure 2C**), the repeated measures ANOVA showed a significant main effect of distance (*F*(1,23) = 71.749, *p* = 1.6e-8), but no significant effect of stimulus noise (*F*(1,23) = 1.041, *p* = 0.3182). The interaction between distance and noise was significant (*F*(2,23) = 6.298, *p* = 0.0196), which was reflected in the steeper decrease of the bias for high compared to low noise stimuli. Post-hoc paired t-tests showed that the response bias was significant for all bins, even for stimuli oriented further away from the boundary (all *p’s* < 0.01).

When considering all trials, instead of only correct trials, we found an overall significant repulsive bias in reproductions away from the discrimination boundary (**Figure 2D**, *F*(1,23) = 5.15, *p* = 0.0329). There was a significant main effect of noise (*F*(1,23) = 37.024, *p* = 3e-6), but no significant effect of distance (*F*(1,23) = 0.205, *p* = 0.65) nor a significant distance x noise interaction (*F*(2,23) = 1.578, *p* = 0.22). Post-hoc t-tests revealed that the bias was pronounced for low noise stimuli (low noise near: *t*(23) = 4.629, *p* = 0.0001; low noise far: *t*(23) = 2.721, *p* = 0.0122), but not for significant for high noise stimuli (high noise near: *t*(23) = 1.635, *p* = 0.12; high noise far: *t*(23) = 0.355, *p* = 0.73). We speculate that the absence of an effect of distance and the lack of bias for high noise stimuli both derive from the fact that boundary judgments about stimuli close to the boundary as well as to high noise stimuli are less strongly coupled to the nominal stimulus orientation, and therefore a systematic effect caused by the boundary judgment is masked in this analysis. Nevertheless, this analysis unequivocally shows that reproduction responses are systematically repelled from the discrimination boundary, even though the boundary is only presented before the stimulus and crucially not during the reproduction phase, thus potentially reflecting a bias in perceptual appearance.

## 3. Experiment 2: Reference repulsion during perception?

In Experiment 1 we found that reproduction responses are systematically repelled from a boundary when people are engaged in a fine discrimination task. Crucially, a stimulus reproduction is necessarily based on a working memory representation of the stimulus which disappeared from view on average two to three seconds ago. Consequently, these reproduction responses are very susceptible to post-perceptual working memory and decision biases. Hence, the bias measured in Experiment 1 could either reflect a bias in the perceived orientation of the stimulus, creating a perceptual illusion, or it may reflect a later working memory or decision-related bias, with the percept of the stimulus being unaffected. In order to test whether reference repulsion is a perceptual illusion, we therefore aimed to probe the perceived orientation of a stimulus more directly in Experiment 2. In particular, the perceived stimulus orientation was measured by asking participants to compare the stimulus to a reference stimulus, simultaneously presented on the screen. This perceptual comparison method reduces the influence of post-perceptual decision and working memory processes (Schneider & Komlos, 2008) as a decision is formed on the basis of material that is available to the observer at the time of the decision. Similar to Experiment 1, we employed stimuli of two noise levels (low/high noise) in order to investigate the influence of stimulus noise on perceptual reference repulsion.

### 3.1. Methods

#### 3.1.1. Participants

Twenty-four new participants took part in Experiment 2. Five participants were excluded before the start of the main experiment, as they performed at chance during the initial practice of the boundary judgment task. The remaining participants (14 female/5 male, age range 18 – 30 years) were naïve to the purpose of the experiment, with the exception of one of the authors. Data of one participant were excluded from data analysis, because no acceptable psychometric model fits could be achieved in multiple conditions. Consequently, data of eighteen participants entered final data analysis, comprising a total of 48,384 trials. All participants reported normal or corrected-to-normal vision and gave written, informed consent prior to the start of the study. The study was approved by the local ethical review board (CMO region Arnhem-Nijmegen, The Netherlands) and was in accordance with the Declaration of Helsinki.

#### 3.1.2. Apparatus and stimuli

The same experimental setup and stimulus parameters as in Experiment 1 were used, except that instead of a response bar at the end of the trial, two orientation stimuli were simultaneously presented on each trial (**Figure 3A**).

**Figure 3:**
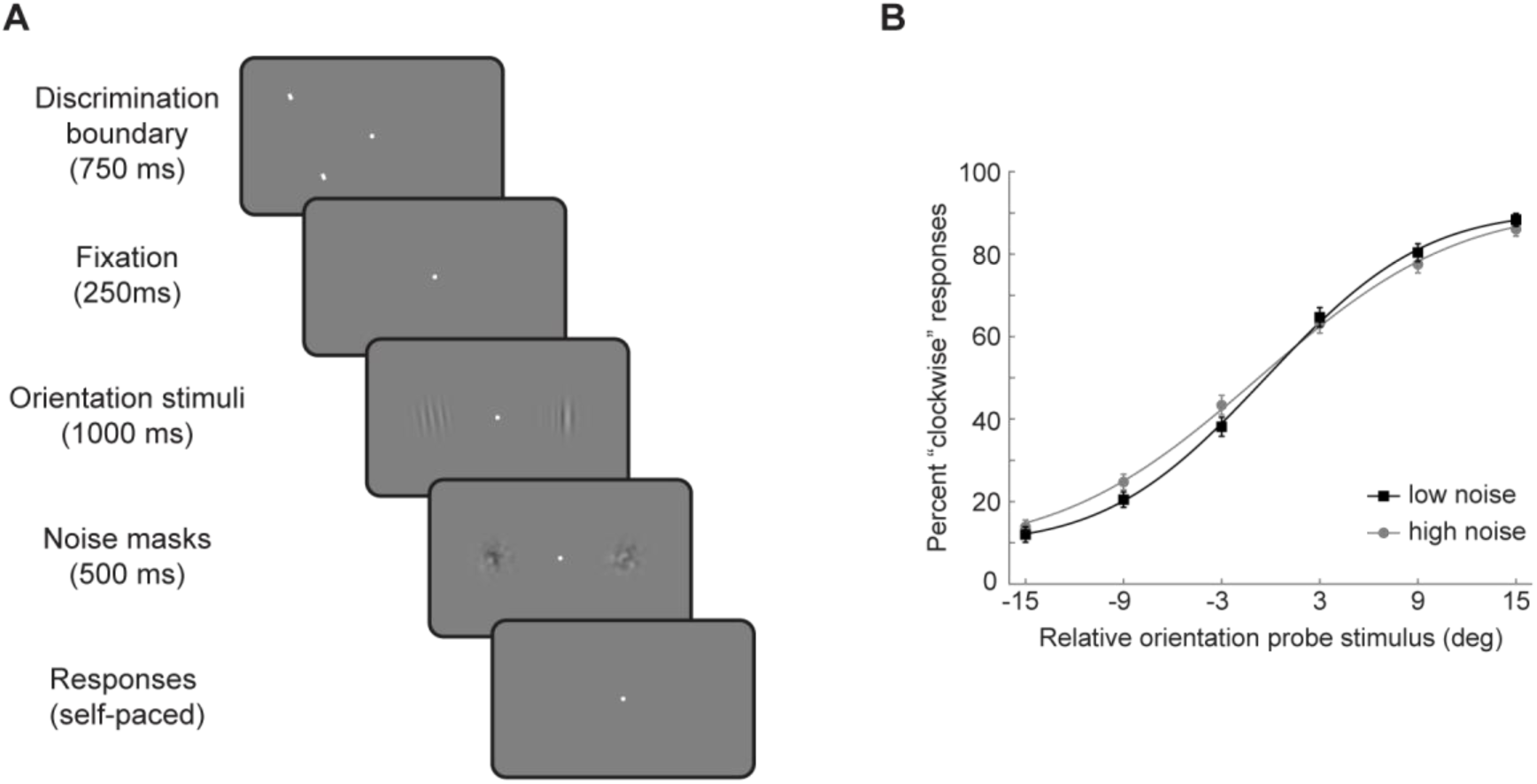
Task and boundary judgment data of Experiment 2 (A) Participants had to indicate whether a probe stimulus was oriented clockwise or counterclockwise with respect to a prior discrimination boundary presented at the same location. Subsequently, participants had to indicate whether the probe stimulus had the same orientation as a reference stimulus simultaneously presented in the opposite visual hemifield. (B) Boundary discrimination judgments of the average observer. Same analysis as in **Figure 2A**. Data points represent group means and error bars depict *SEMs*.

#### 3.1.3. Procedure

At the beginning of each trial an oriented discrimination boundary was presented left or right of fixation for 750 ms. After further 250 ms of fixation, two orientation stimuli of the same noise level were presented simultaneously in the left and right visual field. The stimulus on the side of the prior discrimination boundary (probe stimulus) was oriented 3°, 9° or 15° clockwise or counterclockwise from the discrimination boundary. The stimulus on the opposite side (reference stimulus) had a relative orientation ranging from -12° to 12° in 4° steps with respect to the probe stimulus. After 1000 ms the orientation stimuli were replaced by noise masks, presented for 500 ms. The participants’ task was twofold. First, participants were asked to indicate whether the probe stimulus was oriented clockwise or counterclockwise from the prior discrimination boundary (boundary judgment). Second, participants indicated whether the probe and reference stimuli had the same or a different overall orientation (perceptual comparison judgment). Responses were given after the offset of the masks by successively pressing the left/right arrow key for the boundary judgment and up/down arrow key for the perceptual comparison judgment. The response was followed by a 1-second inter-trial-interval.

Participants completed a total of 2,688 trials in two sessions, each split into 12 blocks. Sessions were conducted on different days, but no more than 5 days apart. The trial sequence was counterbalanced with respect to the horizontal location of the discrimination boundary, noise level of orientation stimuli, probe stimulus orientation with respect to the discrimination boundary and reference stimulus orientation with respect to the probe stimulus. Trials were presented in pseudorandom order. Importantly, within each session, each particular pair of orientation stimuli was presented twice – on one trial the discrimination boundary was oriented clockwise, on the other trial counterclockwise with respect to the probe stimulus. This allowed to study the influence of the boundary judgment on perception with physically exactly matched orientation stimuli. Except for this matching of trials pairs with physically identical orientation stimuli, the overall orientation of the stimuli and discrimination boundaries were randomly drawn from the range of [0, 180°).

At the beginning of the first session, participants practiced the boundary judgment task in isolation in blocks of 48 trials until an acceptable performance level was reached, or were excluded from the experiment if performance was at chance after several blocks. Participants then practiced the main task for at least one block of 56 trials. The practice of the main task was repeated at the beginning of the second session.

#### 3.1.4. Data analysis

Psychometric curves were fit to the boundary judgment data of each individual participant, similar to Experiment 1.

The data of the perceptual comparison judgment were used to measure the perceived orientation of the probe stimulus that was judged with respect to the discrimination boundary. To this end, the probability of a “same” response was expressed as a function of the reference stimulus orientation relative to the probe stimulus. These data were fit with a Gaussian model, with parameter *a, b* and *c*, determining the amplitude, location, and width of the Gaussian, respectively. Importantly, location parameter *b* corresponds to the point of subjective equality (*PSE*), which designates the relative orientation difference between reference and probe stimulus for which participants perceived the orientations as equal. For *b*>0 the reference stimulus had to be oriented more clockwise in order be perceived as equal to the probe stimulus, while for *b*<0 it had to be oriented more counterclockwise. In order to assess whether judging the probe stimulus against the discrimination boundary had a systematic effect on its perceived orientation, the perceptual comparison data were split in two bins according to the relative orientation of the probe stimulus with respect to the discrimination boundary. For this analysis only trials with a correct boundary judgment were taken into account. Hence, the “judgment-cw” bin contained all trials in which participants correctly judged the probe stimulus as more clockwise than the discrimination boundary, whereas the “judgment-ccw” bin contained all trials in which the probe stimulus was correctly judged as more counterclockwise. *PSEs* for ‘judgment-cw’ and ‘judgment-ccw’ bins were separately estimated and the overall bias of the perceived orientation of the probe stimulus was quantified with *ΔPSE* = *(PSE*_*judgment-cw*_ *– PSE*_*judgment-ccw*_) / 2 (**Figure 4A**). For positive *ΔPSEs* the perceived orientation of the probe stimulus was biased away from the discrimination boundary, whereas for negative *ΔPSEs* perceived orientation was biased towards the boundary. *ΔPSEs* were estimated separately for the different absolute orientation differences between probe stimuli and discrimination boundaries, in low and high noise conditions respectively (**Figure 4B, solid lines**). Similar to Experiment 1, a potential caveat here is that participants could be sensitive to random fluctuations in orientation energy of the stochastically generated stimuli. Furthermore, even for physically identical stimuli, the internal representations of those stimuli are likely subject to trial-by-trial fluctuations, due to incomplete sampling of the stimuli and internal noise. Therefore, when considering correct trials only one is prone to only consider those trials in which external and/or internal fluctuation biased the stimulus representation away from the boundary, towards the correct side of the boundary judgment. This would lead to an apparent bias when comparing the probe stimulus to reference stimulus, but would not reflect a systematic bias due to making fine discrimination judgments per se. In order to estimate the contribution of random trial-by-trial fluctuations of stimulus representations to the bias measured in perceptual comparison judgments, we devised an artificial observer model grounded in signal theory (Schneider & Komlos, 2008). With this artificial observer model we simulated behavioral responses for each participant in the task, based on stimulus representations that were subject to random trial-by-trial fluctuations. The amount of trial-by-trial fluctuations (perceptual sensitivity) for each participant was estimated from the empirical data. Crucially, the artificial observer model did not have any in-built reference repulsion bias, i.e. no systematic bias of the probe stimulus orientations due to discrimination judgments. Consequently, analyzing simulated responses of this artificial observer model without a reference repulsion bias allowed us to estimate the contribution of trial-by-trial fluctuations to the bias measured in the perceptual comparison judgment (for details on the artificial observer model see **Supplementary Material**). In other words, the simulated biases of the artificial observer model served as a baseline against which potential reference repulsion biases could be evaluated (**Figure 4b, dotted lines**). In order to statistically assess whether there was a reference repulsion bias, over and above the apparent biases introduced by random trial-by-trial fluctuations in internal/external stimulus representations, we compared the empirical biases with the simulated biases for each boundary distance and noise level using one-sided paired t-tests at a significance level of *α* = 0.05. Furthermore, we also employed one-sided Bayesian paired t-tests, as implemented in JASP, in order to quantify evidence for the null hypothesis of no difference between empirical and simulated biases, against the alternative hypothesis of stronger biases in the empirical data, indicating the presence of a reference repulsion bias.

**Figure 4:**
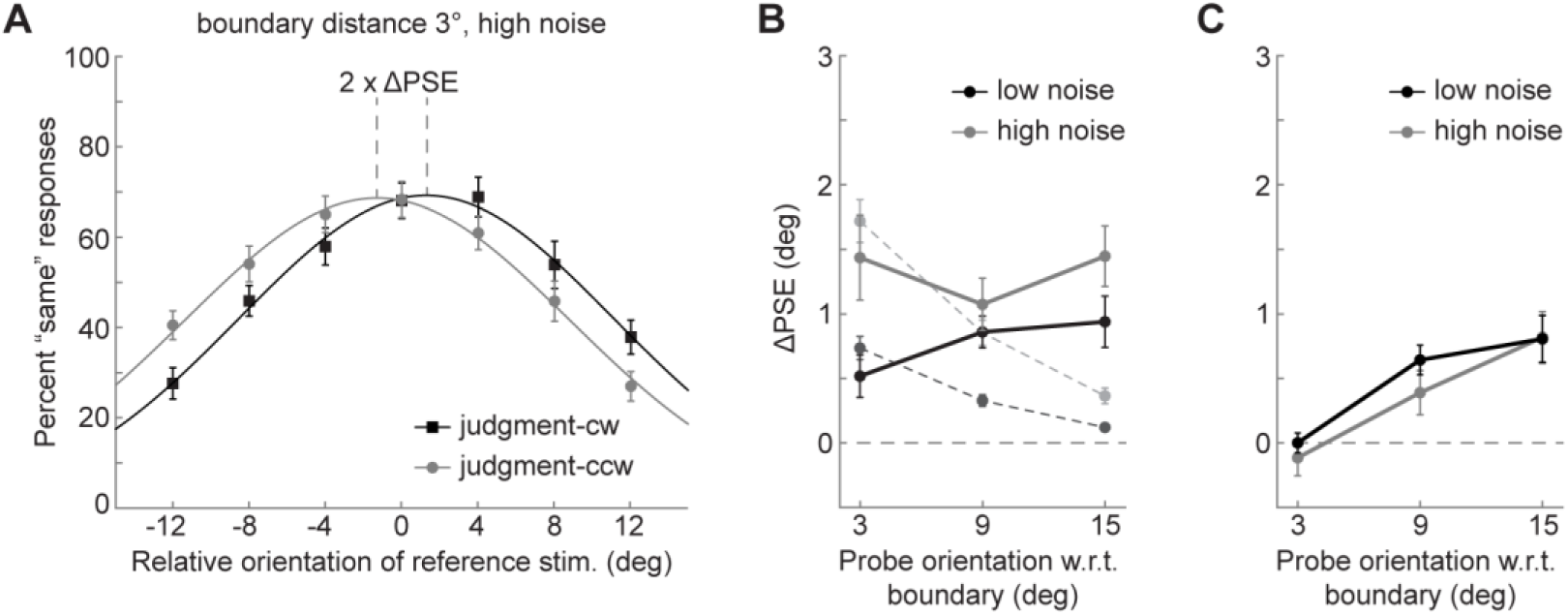
Perceptual comparison between probe and reference stimulus in Experiment 2. (A) Example response distributions of the average observer for the perceptual comparison judgment between probe and reference stimuli containing high noise, when the probe stimulus was oriented 3° from the discrimination boundary. Participants had to indicate whether the probe and reference stimulus had the same or a different orientation. We expressed the probability of a ‘‘same’’ response (*y* axis) as a function of the relative orientation of the reference stimulus with respect to the probe. For positive *x* values, the reference stimulus was oriented more clockwise. Black data points represent trials in which the participant correctly judged the probe stimulus as more clockwise than the discrimination boundary, while grey data points likewise indicate trials with correct counterclockwise boundary judgments. The Gaussian model fits (black and grey lines) indicate that the perceived orientation of the probe stimulus is biased away from the boundary (B) Analysis 1: Group bias for low and high noise trials (black and grey data points, solid lines) for all distances between probe and discrimination boundary (*x* axis). Biases are computed by binning trials according to the boundary judgment for correct trials only (see panel (A)). Data points connected by dotted lines show the biases of the simulated observers without reference repulsion, which serve as a baseline containing biases that are introduced by random trial-by-trial fluctuations in external and internal signals (see **Supplementary Material**). (C) Analysis 2: Same as in (B) but biases are computed by binning trials according to the probe orientation with respect to the discrimination boundary, taking all trials into account. All data points represent group means and error bars depict *SEMs*.

Similar to Experiment 1, an alternative analysis, which avoids the problem of potentially binning according to random trial-by-trial fluctuations, involves taking both correct and incorrect trials into account when binning the data according to the probe stimulus tilt with respect to the discrimination boundary. However, as explained above this analysis can only capture biases related to boundary judgments when boundary judgments are strongly correlated with actual stimulus orientation, e.g. when the probe stimulus is oriented further away from the discrimination boundary. Nevertheless, it presents an informative complement to the first analysis and the comparison to the artificial observer model.

### 3.2. Results

Participants were able to successfully perform the boundary judgment task (**Figure 3B**). They showed a higher sensitivity for low noise stimuli than for high noise stimuli (low noise *β = 0.122 +- 0.010 (SEM),* high noise *β = 0.098 +- 0.005 (SEM); t*(17) = 3.372, *p* = 0.004), whereas there was no significant general bias (low noise *α = - 0.284 +- 0.593 (SEM),* t(17) = - 0.4787, p = 0.6383; high noise *α =- 0.685 +- 0.688 (SEM),* t(17) = - 0.9954, p = 0.3335) and no significant difference across conditions (*t*(17) = 1.219, *p* = 0.24). Further, there was no significant difference in lapse rate (low noise λ = 0.055 +- 0.012, high noise λ = 0.062 +- 0.011; *t*(17) = - 0.486, *p* = 0.63).

When quantifying the perceived orientation of the probe stimulus participants showed a repulsive bias away from the discrimination boundary, in the direction of their (correct) boundary judgment (**Figure 4B, solid lines**, *F*(1,17) = 38.911, *p* = 9e-6). Crucially however, when comparing the bias to the one that was simulated with observers lacking reference repulsion (**Figure 4B, dotted lines**), we found that the biases for probe stimuli close to the discrimination boundary are fully accounted for by random fluctuations in external and/or internal stimulus representations and therefore do not appear to constitute systematic biases caused by the boundary judgment itself (3° low noise *t*(17) = -1.871, *p* = 0.961, BF_+0_. = 0.098; 3° high noise *t*(17) = - 0.812, *p* = 0.786, BF_+0_ = 0.147; 9° high noise *t*(17) = 1.146, *p* = 0.134, BF_+0_ = 0.732). Analyzing trial pairs of physically matched stimuli, we found that the biases for probe stimuli close to the orientation boundary mostly reflected fluctuations in the internal stimulus representations, and only to a very small degree fluctuations in the physical stimuli (see **Supplementary Material**).

The comparison of the results of Experiment 2 with those of Experiment 1 (**Figure 4B** and **Figure 2B**) further supports the hypothesis that the biases found in the reproduction responses do not reflect perceptual, but post-perceptual biases. The biases in reproduction responses of Experiment 1 clearly exceed the apparent biases measured with the perceptual comparison technique in Experiment 2. This difference is particularly evident for stimuli that were tilted 3° from the discrimination boundary (two-sample t-tests, corrected for unequal variances where necessary; low noise: *t*(28.066) = 6.154, *p* = 1e-6; high noise: *t*(40) = 4.127, *p* = 1.81e-6), significant for low but not for high noise stimuli tilted 9° from the boundary (low noise: *t*(25.596) = 3.446, *p* = 0.002; high noise: *t*(31.326) = 1.684, *p* = 0.1021), completely absent for low noise stimuli tilted 15° from the boundary (*t*(28.222) = 1.514, *p* = 0.14) and only opposite for 15° high noise stimuli (*t*(29.874) = 0.-2.217, *p* = 0.034). Furthermore, assuming that the influence of the random trial-by-trial fluctuations in the stimulus representations were approximately equal across the two experiments, the additional bias in reproduction responses shown in **Figure 2B** over the bias in perceptual comparisons shown in **Figure 4B** cannot be explained by random-trial-by-fluctuations and thus appears to reflect a genuine post-perceptual bias in reproduction responses, which decreases with increasing orientation difference between stimulus and boundary.

Importantly, when comparing the empirical biases of Experiment 2 with the simulated observer lacking reference repulsion (**Figure 4B**), probe stimuli oriented further away from the discrimination boundary, at 9° (low noise: *t*(17) = 4.696, *p* < 0.001, BF_+0_ = 294.898) and 15° (low noise: *t*(17) = 4.043, *p* < 0.001, BF_+0_ = 86.545; high noise: *t*(17) = 4.392, *p* < 0.001, BF_+0_ = 166.905) orientation difference, could not be explained by random trial-by-trial fluctuations and seem to reflect a genuine systematic repulsive perceptual bias, away from the discrimination boundary, in the direction of the boundary judgment.

In order to confirm that the biases for probe stimuli oriented further away from the discrimination boundary could not be explained by random trial-by-trial fluctuations in stimulus representations we conducted a second analysis in which we binned trials according to the probe stimulus tilt with respect to the decision boundary, taking both correct and incorrect trials into account. As this analysis did not factor in the participants’ boundary judgments, any bias observed here would indicate the presence of a systematic perceptual bias related to the discrimination boundary. Indeed, we observed a significant bias away from the boundary (**Figure 4c**, *F*(1,17) = 15.342, *p* = 0.001). The bias increased with increasing orientation difference between stimulus and boundary (*F*(2,34) = 30.398, *p* = 2.7 x 10e-8). We did not observe a significant difference across noise levels (*F*(1,17) = 1.790, *p* = 0.199). Neither was there a significant interaction between noise level and boundary distance (*F*(2,34) = 0.417, *p* = 0.662). Post-hoc two-sided one-sample t-tests revealed that the bias was present for stimuli oriented 9° and 15° from the decision boundary (9° low noise: *t*(17) = 5.448, *p* < 0.001, high noise: *t*(17) = 2.207, *p* = 0.041; 15° low noise: *t*(17) = 4.295, *p* < 0.001, high noise: *t*(17) = 3.963, *p* = 0.001), but not for stimuli oriented 3° away from the decision boundary (3° low noise: *t*(17) = -6e-4, *p* = 0.99; 3° high noise: *t*(17) = - 0.8, *p* = 0.435). These results are congruent with the results of the first analysis in indicating that there is no perceptual bias for stimuli oriented close to the discrimination boundary, but a genuine perceptual bias for probe stimuli oriented further away from the boundary. In particular, the perceived orientations of these probe stimuli are systematically biased away from the boundary.

## 4. Experiment 3: Perceptual bias due to discrimination judgments?

Although in Experiment 2 there was no perceptual reference repulsion for stimuli oriented close to the discrimination boundary, the perceived orientation of stimuli oriented further away from the boundary was repelled. This stands in stark contrast to previous findings of reference repulsion, where the strongest bias for stimuli oriented was observed close to the discrimination boundary (Jazayeri & Movshon 2007; Luu & Stocker, 2018; see also Experiment 1 in this study - for comparisons see **Supplemental Material**). In fact, the tuning of the perceptual bias in Experiment 2 appears more congruent with classical negative adaptation effects, for which maximal repulsion is typically found for orientation differences of around 20° (Gibson & Radner, 1937). This begs the question whether the perceptual bias we found in Experiment 2 is different in nature to reference repulsion. Importantly, theories of reference repulsion postulate that the bias is dependent on judging a stimulus against a discrimination boundary or reference. Conversely, reference repulsion should not occur if one is not engaged in a fine discrimination task against a boundary. In line with this, Jazayeri and Movshon (2007) found that reproduction responses were only biased away from the boundary when observers were engaged in a fine discrimination task, whereas responses were biased towards the boundary when observers performed a coarse discrimination task. Besides differences in orientation tuning, this distinguishes reference repulsion from classical adaptation effects for which no comparison between an adaptor and test stimulus is required. Therefore, in Experiment 3 we tested whether the perceptual bias of Experiment 2 would vanish when participants were not explicitly engaged in a fine boundary discrimination task.

### 4.1. Methods

#### 4.1.1. Participants

Twelve naïve participants took part in Experiment 3 (10 female, age range 19 – 30 years), providing a total of 24,000 trials. The sample size was motivated by a power analysis based on the effect size found in Experiment 2, achieving 90% power for detecting a repulsive bias for low noise stimuli oriented 15° from the boundary. None of these participants took part in Experiment 1 or 2. All participants reported normal or corrected-to-normal vision and gave written, informed consent prior to the start of the study. The study was approved by the local ethical review board (CMO region Arnhem-Nijmegen, The Netherlands) and was in accordance with the Declaration of Helsinki.

#### 4.1.2. Apparatus and stimuli

Experimental setup and stimuli were similar to Experiment 2. In contrast to Experiment 2 only low noise stimuli (*κ* = 500) were used. Furthermore, on a subset of trials, instead of the orientation stimuli, a white response bar stimulus (length 4°, width 0.09°) was presented at 10° eccentricity after the offset of the boundary, prompting participants to reproduce the orientation of the boundary.

#### 4.1.3. Procedure

Similar to Experiment 2, at the beginning of each trial an oriented boundary was presented left or right of fixation for 750 ms (**Figure 5**). The orientation of the discrimination boundary was randomly drawn from the range of all possible orientations, [0-180°). On 2/3 of the trials, two orientation stimuli were presented simultaneously in the left and right visual field, 250ms after the offset of the boundary. The stimulus on the side of the prior boundary (probe stimulus) was oriented 3°, 9° or 15° clockwise or counterclockwise from the boundary. The stimulus on the opposite side (reference stimulus) had a relative orientation ranging from -12° to 12° in 4° steps with respect to the probe stimulus. After 1000ms the orientation stimuli were replaced by noise masks, presented for 500 ms. On those perceptual comparison trials, participants indicated whether the two orientation stimuli had the same or a different overall orientation. Responses were given after the offset of the masks by pressing the up/down arrow keys. The response was followed by a 1-second inter-trial-interval. Importantly, in contrast to Experiment 2 participants never had to compare the orientation stimuli to the boundary. In order to ensure that participants still attended to the boundary, on 1/3 of the trials a white response bar was presented in the hemifield of the boundary 250ms after its disappearance. On those boundary reproduction trials, the participant had to reproduce the orientation of the boundary by adjusting the response bar with the left and right arrow key. The response was submitted by pressing the space bar. The response was followed by a 1-second inter-trial-interval, before the next trial began.

**Figure 5:**
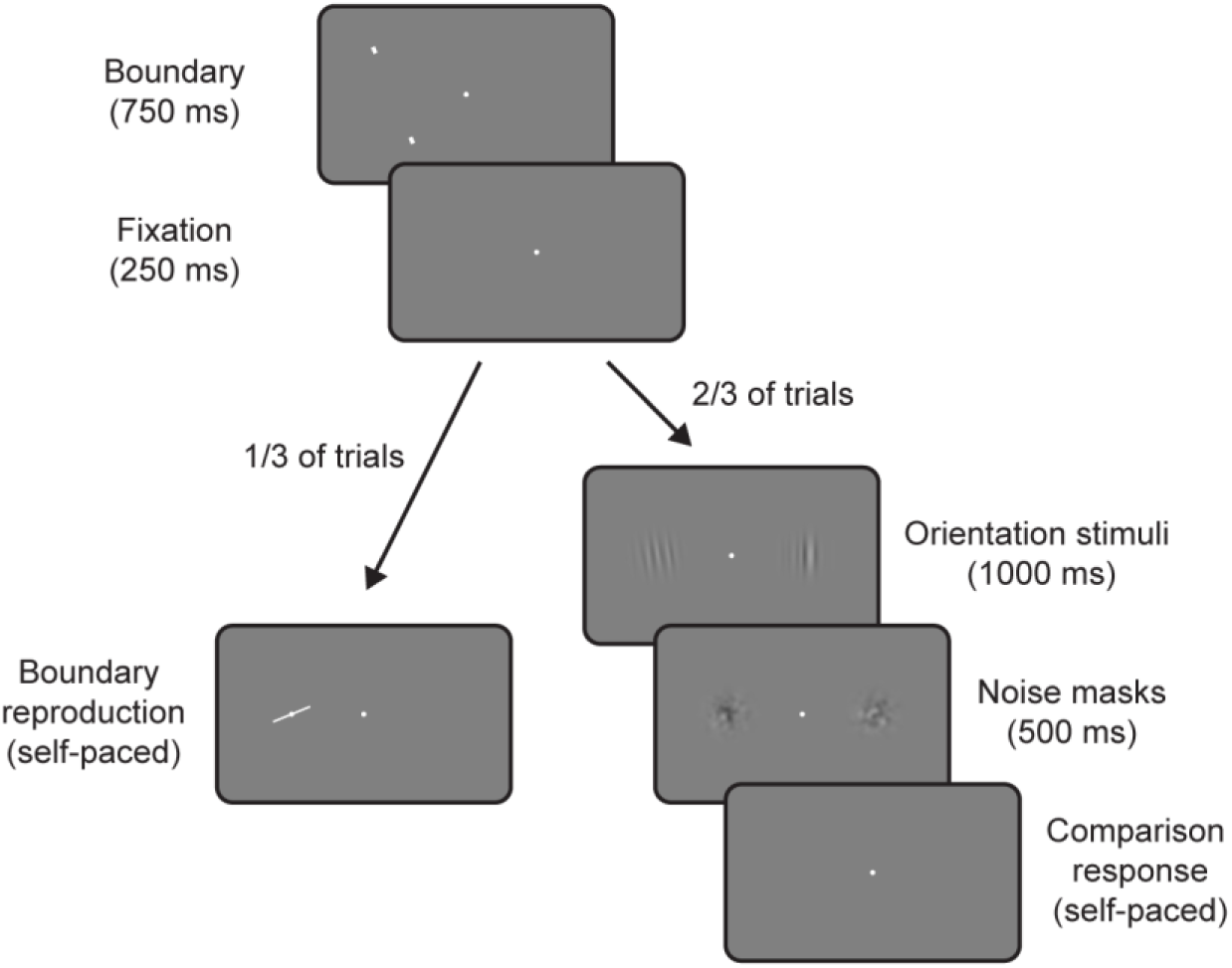
Trial design of Experiment 3. On 2/3 of trials participants indicated whether two simultaneously presented orientation stimuli had the same or a different overall orientation (perceptual comparison trials). On 1/3 of randomly interleaved trials, instead of the orientation stimuli a white response bar appeared, prompting participants to adjust it to the orientation of the prior boundary (boundary reproduction trials).

Participants completed a total of 2,000 trials in two sessions, each split into 10 blocks. Sessions were conducted on different days, but no more than 7 days apart. The trial sequence was counterbalanced with respect to the horizontal location of the boundary, probe stimulus orientation with respect to the boundary and reference stimulus orientation with respect to the probe stimulus. Trials were presented in pseudorandom order and boundary reproduction trials and perceptual comparison trials were randomly interleaved. At the beginning of the first session, participants practiced the main task for at least one block of 56 trials. The practice of the main task was repeated at the beginning of the second session.

#### 4.1.4. Data analysis

The perceptual comparison trials were analyzed similarly to Experiment 2, taking all trials into account. That is, for each orientation difference between probe stimulus and boundary we split the trials into two bins – one bin containing those trials in which the probe stimulus was oriented clockwise from the boundary and one containing those in which it was oriented counterclockwise. For each bin, the probability of a “same” response was expressed as a function of the reference stimulus orientation relative to the probe stimulus. As for Experiment 2, these data were fit with a Gaussian model for which location parameter *b* corresponds to the point of subjective equality (*PSE*), which designates the relative orientation difference between reference and probe stimulus for which participants perceived the orientations as equal. The overall bias of the perceived orientation of the probe stimulus was quantified with *ΔPSE* = (*PSE*_*probe-cw*_ *– PSE*_*probe-ccw*_*) / 2*, separately for each orientation difference between probe stimulus and boundary.

In order to statistically assess whether the perceived orientation of the probe stimulus was biased away from the boundary, even though participants did not explicitly judge the probe stimulus against the boundary, we conducted a repeated measures ANOVA with the orientation difference between the probe stimulus and boundary as the repeated measures factor. Furthermore, in order to test whether the biases in Experiment 3 would be similar in magnitude to the biases in Experiment 2, we conducted a Bayesian repeated measures ANOVA with the orientation difference between the probe stimulus and boundary as the repeated measures factor and “Experiment” as a between subject factor.

### 4.2. Results

Participants were able to accurately reproduce the orientation of the boundary on boundary reproduction trials (mean absolute error 5.44° +- 0.31 SEM; mean standard deviation 7.66° +- 0.48 SEM; mean kurtosis 19.61 +- 4.59), indicating that they successfully attended to the orientation of the boundary. For the perceptual comparison trials, we measured a pronounced repulsive bias of the perceived probe stimulus orientation away from the boundary (**Figure 6**, *F*(1,11) = 25.549, *p* < 0.001). As in Experiment 2, this bias increased with increasing orientation difference between stimulus and boundary (*F*(2,22) = 4.954, *p* = 0.017). Post-hoc one sample t tests revealed a significant bias for 9 and 15° orientation differences (9°: *t*(11) = 2.915, *p* = 0.014; 15°: *t*(11) = 5.124, *p* < 0.001), but not for 3° orientation differences (*t*(11) = 2.024, *p* = 0.068).

**Figure 6:**
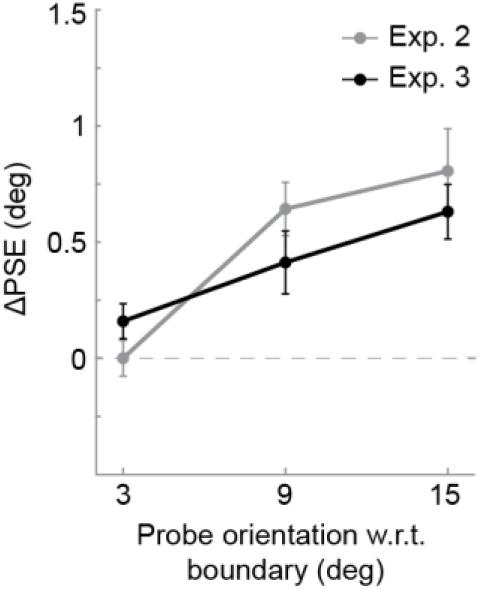
Results of Experiment 3 in which no explicit boundary judgments were made. Group biases for all distances between probe stimulus and boundary (*x* axis). Biases are computed by binning trials according to the probe orientation with respect to the boundary, and computing differences in *PSEs* between the bins. Results of Experiment 2 are replotted in grey (same as in **Figure 4C**, low noise condition). There appears to be no difference in bias magnitude, when the probe stimulus is judged to the discrimination boundary compared to when no such judgment is made. All data points represent group means and error bars depict *SEMs*.

Subsequently, we statistically compared the magnitude of the repulsive bias to the one found in Experiment 2 using a Bayesian repeated measures ANOVA. Compared to the model containing the repeated measures factor of the orientation difference between probe stimulus and boundary, which performed best compared to the null model (BF _boundary distance vs. null_ = 76340.54), there was moderate evidence against additionally including a between subject “Experiment” factor (BF _full main effects model vs boundary distance_ = 0.367) or the interaction between boundary distance and experiment (BF _full model vs boundary distance_ = 0.225). This indicates that the biases measured in Experiment 2 and 3 were of similar magnitude. Consequently, the perceptual bias measured in Experiment 2 does not appear to occur due to judging the probe stimulus against the discrimination boundary and therefore does not reflect a reference repulsion bias as previously described. Rather, the perceptual bias seems to occur due to the encoding of the discrimination boundary per se and resembles classical tilt-aftereffects.

## 5. Discussion

Biases in perceptual decision-making can arise at different stages, influencing perceptual appearance or post-perceptual decision-related processes. The stage at which a bias arises determines what information the organism can utilize at any given point in time, and thus it is crucial to establish the level at which a bias operates. Reference repulsion, a bias due to making fine discrimination judgments, has previously been argued to reflect a bias on perceptual appearance, i.e. a perceptual illusion (Jazayeri & Movshon, 2007; Stocker & Simoncelli, 2008). A recent study by Zamboni et al. (2016) called the perceptual nature of reference repulsion into question, demonstrating that the bias measured with stimulus reproductions crucially depended on task parameters of the post-perceptual reproduction phase. In particular, reference repulsion was only evident when the reference was present during the reproduction response. Although this study indicated that the large biases measured in previous studies were decision-related, it remained an open question whether discrimination judgments could also bias the perceptual appearance of a stimulus. In Experiment 1, we showed that reproduction responses were repelled from a discrimination boundary even when the boundary was only presented prior to a stimulus, and thus not during the reproduction phase. In order to test whether this reflects a perceptual bias, in Experiment 2 we measured perceptual appearance with a more direct and sensitive perceptual comparison technique while participants were engaged in a boundary discrimination task. Although we did find a genuine perceptual bias for stimuli oriented further away from the discrimination boundary, this effect had a very different tuning profile than the reference repulsion bias in reproduction responses, and did not necessitate an explicit discrimination judgment of the biased stimulus against the boundary (Experiment 3). In terms of orientation tuning, the perceptual bias rather resembled a tilt illusion (Clifford, 2014; Gibson, 1937; Schwartz, Hsu & Dayan, 2007; Wenderoth & Johnstone, 1987) or tilt-aftereffect (Gibson & Radner, 1937; Webster, 2015). Consequently, our experiments provide evidence against the perceptual nature of reference repulsion, indicating that making fine discrimination judgments does not alter the appearance of visual stimuli, but instead leads to a post-perceptual working memory or decision bias.

In Experiment 1, we found that reproduction responses were systematically biased away from the discrimination boundary, even though the boundary was not present during the reproduction phase. This result appears to contradict a previous study, which did not find reference repulsion when the discrimination boundary was absent during the reproduction phase (Zamboni et al., 2016; but see reanalysis by Luu & Stocker, 2018). However, when analyzing the current data in a similar way as Zamboni et al., that is quantifying the bimodality of the response distributions, only one of twenty-four participants showed significantly bimodal response distributions in the current experiment (for details see **Supplemental Material**). Furthermore, the response histograms of the average observer did not show any clear signs of bimodality (see **Supplemental Material**), in line with the findings by Zamboni et al. This is likely because the current reference repulsion bias, albeit clear and statistically significant, was too subtle to lead to clearly bimodal response distributions and emphasizes the importance of analyzing the data in a different way to detect and quantify these subtle reference repulsion biases. Nevertheless, given the much smaller magnitude of the bias compared to previous studies which presented the reference during the reproduction phase (Jazayeri & Movshon, 2007; Luu & Stocker, 2018; Zamboni et al., 2016), the current results are in line with the observation by Zamboni et al. that the reference repulsion bias is greatly reduced when removing the reference during reproduction. These findings beg the question whether the smaller bias, occurring when the reference was absent during reproduction, reflects a perceptual bias, which would exist alongside a stronger post-perceptual bias that is dependent on the presentation of the reference during the reproduction phase. Therefore, the perceptual nature of the reference repulsion bias was investigated in Experiment 2.

In order to minimize the influence of post-perceptual working memory and decision processes, which could have played a role in the reproduction responses of Experiment 1, in Experiment 2 we measured the perceived orientation of a stimulus more directly, in comparison to a simultaneously presented reference stimulus. Such a direct perceptual comparison minimizes the influence of working memory and post-perceptual decisions (Fritsche et al., 2017; Schneider & Komlos, 2008) and thus could shed light on the perceptual nature of the reference repulsion bias. Notably, even though participants only communicated their decision about the perceptual comparison after the stimuli had disappeared from the screen, and therefore had to store their decision in working memory, a binary decision about the equality of two stimuli is more robust to fluctuations and biases in working memory, compared to a representation of a continuous variable such as the overall orientation of a grating stimulus. To investigate whether discrimination judgments would alter the perceived orientation of a stimulus we computed biases between bins of trials with opposite and correct boundary judgments. However, taking only correct trials into account has the side effect of potentially making random fluctuations in stimulus representations systematic. That is, even before a discrimination judgment is made, the underlying stimulus representation might be randomly biased due to external or internal noise. Since discrimination judgments should be, to some degree, sensitive to these random biases, taking only correct trials into account can lead to a biased selection of trials, predominantly retaining those trials in which the stimulus representation was biased in favor of the correct boundary judgment. In order to quantify the contribution of these random trial-to-trial fluctuations to our reference repulsion estimates we simulated artificial observers without reference repulsion. The simulations revealed that variability of stimulus representations contributed strongly to the estimated reference repulsion effect when stimuli were oriented close to the discrimination boundary. In fact, the empirically estimated reference repulsion effect for stimuli close to the discrimination boundary was fully explained by random trial-by-trial fluctuations and therefore did not constitute a systematic bias due to making fine discrimination judgments. Importantly, with the help of further simulations we confirmed that our analysis was in principle able to detect even very small reference repulsion biases of 0.2 to 2 degrees if they would have been present in the data (see **Supplemental Material**), further underlining the sensitivity of the perceptual comparison paradigm. It is important to note, however, that the ability to detect reference repulsion effects in the perceptual comparison task depends on two conditions. First, participants need to make the discrimination judgments before the perceptual comparison judgments, so that discrimination judgments could in principle bias the appearance of the probe stimuli. If, however, participants would make the judgments in the opposite order, no reference repulsion could be detected in the perceptual comparison data. Although we were unable to experimentally impose a definite order of judgments, participants were explicitly instructed to make the judgments in the intended order. Furthermore, since the discrimination boundary disappeared before the onset of the orientation stimuli and thus had to be encoded in working memory, it appears more efficient to make the judgment involving the boundary information first. A control experiment, in which we varied the instructed order of judgments, confirmed that our instructions about the order of judgments were likely effective and that making the boundary judgment first presented the more efficient order, leading to a markedly higher task performance that was comparable to Experiment 2 (for details see **Supplemental Material**). Therefore, participants likely adhered to the instructed order of judgments during Experiment 2. A second necessary condition to measure reference repulsion is that the effect should be to some degree spatially specific. If reference repulsion would be completely spatially unspecific the bias would spread to both orientation stimuli and bias both stimuli to an equal amount. By consequence, the comparison between the two orientation stimuli would then be unbiased. In order to minimize a spreading of perceptual biases to both orientation stimuli we presented the stimuli 20 visual degrees apart. However, although very unlikely for perceptual biases, we cannot completely rule out the possibility that perceptual reference repulsion without any spatial specificity does exist.

The presence of reference repulsion in reproduction responses of Experiment 1 and the absence of a reference repulsion bias in perceptual comparison responses of Experiment 2 point towards a post-perceptual decision or working memory locus of the bias. Congruent with a working memory bias, a recent study found repulsive interactions of orientations held in working memory with strongest biases for similar orientations (Bae & Luck, 2017). This effect could be explained by a relational representation model in which each item serves as a reference for representing the other item. It appears plausible that a similar repulsive working memory bias could occur when representing a memorized oriented stimulus to an external reference boundary. Related to this, a recent study proposed a Bayesian decoding model from high-to low-level features as the source of the repulsive bias between successively reproduced line orientations and reference repulsion (Ding, Cueva, Tsodyks & Qian, 2017). Although highlighting the role of working memory in their model, Ding et al. claimed that the repulsive biases reflect a perceptual decoding strategy. Crucially, in the current study we show that when reducing the working memory load between stimulus presentation and response to a minimum there are no such repulsive decoding biases in perception (Experiment 2). In other words, while decoding sensory representations for perception is initially unbiased, later decoding of working memory representations could be biased in a repulsive manner. Therefore, rather than explaining reference repulsion and repulsive biases between working memory items as “misperceiving” visual features due to perceptual decoding strategies, we view these biases as potentially “misremembering” previously perceived visual features, perhaps due to biased working memory decoding strategies. Another recent theory explains reference repulsion as the result of a self-consistency principle in perceptual inference (Luu & Stocker, 2018). According to this theory, estimations of a stimulus feature are not only conditioned on the stimulus information, but also on the participant’s preceding discrimination judgment, in order for discrimination judgments and estimations to be consistent with each other. Our results are broadly in line with this theory, but indicate that the representation that is used for the perceptual comparison in Experiment 2, which likely underlies the percept of the stimulus, is not conditioned on the preceding discrimination judgment. It rather seems, that consistency would be achieved by conditioning a working memory or higher-level decision-related representation of the stimulus, without affecting the perceptual appearance of the stimulus.

In order to further advance our understanding of the post-perceptual reference repulsion it may be fruitful for future studies to investigate which post-perceptual factors can influence reference repulsion in reproduction responses. For instance, it is conceivable that the reference repulsion bias could be modulated by the temporal delay between stimulus presentation and reproduction, similar to attractive effects exerted by the stimulus history (Akrami, Kopec, Diamond & Brody, 2018; Bliss et al., 2017; Fritsche et al., 2017; Papadimitriou et al. 2015). Furthermore, the post-perceptual nature of this repulsive bias away from an external reference delineates the effect from repulsive perceptual biases away from cardinal orientations, which may act as internal references (Rauber & Treue, 1998; Tomassini, Morgan & Solomon, 2010; Wei & Stocker, 2015). It thus seems that repulsive biases can originate at multiple stages during perceptual decision making and potentially jointly affect perceptual decisions.

In contrast to the lack of bias for stimuli oriented close to the discrimination boundary, we did find a genuine perceptual bias for stimuli oriented further away from the boundary. In particular, the perceived orientation of a probe stimulus was biased away from the boundary, and this bias increased with increasing orientation difference between probe stimulus and boundary. Importantly, this orientation tuning was very different compared to previous reports of reference repulsion, for which the biases decreased with increasing orientation difference between stimulus and boundary. Furthermore, in Experiment 3 we found that explicit discrimination judgments are not necessary for this bias to occur. We interpret this as further evidence that the observed perceptual bias does not reflect reference repulsion, for which making fine discrimination judgments is necessary (Jazayeri & Movshon, 2007). However, even though Experiment 3 measured the perceptual appearance of a stimulus when participants were not required to compare the stimulus to a discrimination boundary, it is nevertheless possible that participants still made such a discrimination judgment, explicitly or implicitly. As no such comparison was required by the task, this seems rather unlikely. If anything, participants could have used the boundary orientation as a cue about the rough orientation of the upcoming orientation stimuli. However, according to Jazayeri and Movshon’s theory of reference repulsion, such a coarse comparison should have led to attractive rather than repulsive biases (see Experiment 2 in Jazayeri & Movshon, 2007). Nevertheless, it should be mentioned that a previous study found systematic reference repulsion biases in direction estimates even when observers were not required to make an explicit prior discrimination judgment (Experiment 2 in Zamboni et al., 2016). Importantly, in contrast to our Experiment 3, participants had to reproduce motion direction from memory and the discrimination boundary was presented during the entire trial. For such an experimental design, it seems plausible that participants used the discrimination boundary as an anchor for their fading memory representation of the stimulus in order to optimize their performance for the direction reproductions. Thus, participants may have been encouraged by the task design to make a discrimination judgment, even though this was not explicitly asked for. Conversely, in our Experiment 3, the orientation of the boundary was not informative about the differences between the orientation stimuli that were important for the same/difference judgment. Thus, we believe that, in contrast to Experiment 2 by Zamboni et al., the incentive to compare the orientation stimuli to the boundary was minimal in our experimental design. Instead we offer two alternative explanations for the repulsive perceptual biases found in Experiment 2 & 3. First, since the discrimination boundary was presented in the form of small line segments that were not overlapping in space or time with the presentation of the orientation stimuli, it is conceivable that participants represented an imagined orientation boundary connecting the two line segments in order two solve both boundary tasks of Experiment 2 & 3. It has been previously reported that negative after-effects can occur due to mentally generated lines (Mohr, Linder, Dennis, Sireteanu, 2011) and orientation stimuli encoded in working memory (Saad & Silvanto, 2013; Scocchia, Cicchini & Triesch, 2013). Therefore, the current negative aftereffect might be similarly caused by briefly mentally generated internal representations of the discrimination boundary. Notably, in our experiments such an adaptation effect would develop over very short timescales of a few hundred milliseconds, which would extend previous findings that used much longer adaptation periods. Second, it is possible that the bias reflects a tilt illusion induced by the line segments of the discrimination boundary acting as a surrounding context to the probe stimulus. Although the presentations of the boundary and probe stimulus were separated by 250 ms, it has been reported that the tilt illusion can persist over short stimulus onset asynchronies between center and surround stimuli (Corbett, Handy & Enns, 2009). However, it must be noted that in our experiments the spatial frequencies of the orientation stimuli and the discrimination boundary were very different and the magnitude of the tilt illusion was previously found to decrease with increasing differences in spatial frequency between center and surround stimulus (Georgeson, 1973). In order to investigate whether the perceptual bias observed in the current study is due to a tilt-aftereffect to mentally generated lines, or reflects a tilt-illusion in response to a previously presented orientation boundary, future studies could use orientation boundaries without local orientation information, e.g. two opposing dots instead of two opposing line segments, which should eliminate a tilt-illusion effect to an oriented surround.

## 6. Conclusion

Our results demonstrate that discriminating a stimulus against an external reference does not lead to a repulsive perceptual bias away from the reference. This suggests that reference repulsion measured in the current and previous studies reflects a post-perceptual decision or working memory related phenomenon and does not constitute a perceptual illusion. The finding underlines the importance of studying and separating the different stages at which biases in perceptual decision making can occur.

## Supplementary material

Data and code are available from the Donders Institute for Brain, Cognition and Behavior repository at http://hdl.handle.net/11633/di.dccn.DSC_3018029.03_140.

## Acknowledgements

We thank Micha Heilbron and Alexis Pérez Bellido for comments on a previous version of the manuscript. M.F. was supported by a grant from the European Union Horizon 2020 Program (ERC Starting Grant 678286, “Contextvision”). F.P.d.L. was supported by grants from NWO (VIDI grant 452-13-016), the James S McDonnell Foundation (Understanding Human Cognition, 220020373) and the European Union Horizon 2020 Program (ERC Starting Grant 678286, “Contextvision”).

Commercial relationships: none

## Artificial observer model

### 1. Signal theory

For the discrimination judgment of the probe stimulus orientation against the discrimination boundary, the probability for judging the probe stimulus as clockwise from the boundary is given by

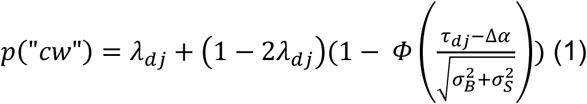

where Δ*α* denotes the signed orientation difference between the probe stimulus and discrimination boundary, *τ*_*dj*_ is the criterion above which the observer will report that the probe stimulus is more clockwise than the boundary (*τ*_*dj*_ = 0 for an unbiased observer), 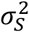 is the variance in the stimulus sampling distribution, 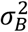 is the variance in the discrimination boundary sampling distribution, *λ*_*dj*_ is the lapse rate and *Φ* denotes the cumulative normal distribution.

For the perceptual comparison judgment between probe and reference stimulus, the probability for judging the two stimuli as having the same orientation is given by

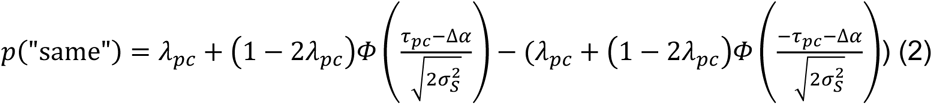

where Δ*α* denotes the signed orientation difference between the probe and reference stimulus, [−*τ*_*pc*_, *τ*_*pc*_] determines the range of orientation differences for which the observer will report the two stimuli having the same orientation and 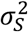 denotes the variance in the stimulus sampling distributions of probe and reference stimuli, respectively. Note, that we assume that estimates of probe and reference orientation are sampled from distributions of equal variance. *λ*_*pc*_ denotes the lapse rate in the perceptual comparison (equality) judgment. For more information on the underlying signal theory of comparative and equality judgments see Schneider & Komlos (2008).

### 2. Estimating parameters for individual participants

We estimated variance, criterion and lapse parameters for each individual participant from the empirical data. In particular, we estimated variance 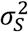 for probe and reference stimuli as well as criterion *τ*_*pc*_ and *λ*_*pc*_ from each participant’s perceptual comparison judgments, separately for low and high noise stimuli, by minimizing the squared error between the empirical data and model predictions. Subsequently, we estimated the variance of the boundary sampling distribution 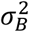, criterion *τ*_*dj*_ and *λ*_*dj*_ from the discrimination judgment data, by jointly minimizing the squared error for discrimination judgments of low and high noise stimuli. While 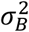 was constrained to be equal for low and high noise trials, *τ*_*dj*_ and *λ*_*dj*_ was allowed to be different across noise levels. Parameter estimates for the participants of Experiment 2 are shown in **Table T1**.

**Table T1.** Parameter estimates for participants of Experiment 2 (see sections 1 & 2).

| Subject ID | $\sigma_s$ (low / high noise) | $\sigma_B$ | $\tau_{dj}$ (low / high noise) | $\tau_{pc}$ (low / high noise) | $\lambda_{dj}$ (low / high noise) | $\lambda_{pc}$ (low noise / high noise) |
| --- | --- | --- | --- | --- | --- | --- |
| 1 | 2.25 / 3.35 | 8.56 | 1.64 / 0.22 | 5.25 / 11.28 | 0.04 / 0.04 | ~ 0 / 0.09 |
| 3 | 3.00 / 7.10 | 15.45 | -1.13 / 1.08 | 6.54 / 11.30 | 0.09 / 0.09 | 0.12 / ~ 0 |
| 5 | 3.18 / 5.92 | 6.23 | 0.59 / -1.56 | 5.25 / 8.10 | 0.08 / 0.08 | ~ 0 / ~ 0 |
| 6 | 2.79 / 5.65 | 9.70 | -0.23 / -0.83 | 6.96 / 13.87 | 0.15 / 0.15 | 0.14 / 0.21 |
| 7 | 3.12 / 3.72 | 6.08 | -0.83 / -0.93 | 5.19 / 10.24 | 0.03 / 0.03 | ~ 0 / 0.12 |
| 8 | 2.93 / 6.22 | 6.27 | 0.61 / 0.08 | 3.68 / 9.75 | 0.07 / 0.07 | ~ 0 / 0.01 |
| 9 | 3.27 / 5.60 | 14.76 | 4.26 / 0.33 | 4.92 / 6.58 | ~ 0 / ~ 0 | ~ 0 / ~ 0 |
| 11 | 4.50 / 7.03 | 9.59 | 2.65 / -0.36 | 6.03 / 10.14 | 0.08 / 0.08 | ~ 0 / ~ 0 |
| 12 | 3.17 / 6.77 | 13.15 | 0.04 / -0.24 | 5.76 / 7.04 | 0.07 / 0.07 | ~ 0 / ~ 0 |
| 14 | 2.30 / 4.56 | 4.31 | 1.02 / -1.24 | 3.91 / 8.70 | 0.10 / 0.10 | ~ 0 / ~ 0 |
| 15 | 3.19 / 2.30 | 12.06 | -4.43 / 3.43 | 3.11 / 10.42 | 0.04 / 0.04 | ~ 0 / 0.33 |
| 17 | 3.23 / 3.92 | 7.23 | -5.07 / -4.74 | 7.17 / 12.32 | 0.07 / 0.07 | ~ 0 / 0.06 |
| 18 | 3.26 / 4.69 | 8.31 | -0.02 / -1.73 | 6.73 / 11.64 | 0.14 / 0.14 | ~ 0 / 0.02 |
| 19 | 2.84 / 5.56 | 6.46 | -0.42 / 0.36 | 3.70 / 3.91 | 0.09 / 0.09 | ~ 0 / ~ 0 |
| 21 | 5.29 / 4.03 | 6.86 | -2.81 / -2.41 | 8.20 / 13.96 | ~ 0 / ~ 0 | ~ 0 / 0.11 |
| 22 | 2.84 / 4.08 | 5.26 | 1.79 / 2.61 | 3.70 / 8.11 | 0.07 / 0.07 | ~ 0 / ~ 0 |
| 23 | 3.35 / 6.18 | 4.01 | -3.51 / -5.80 | 4.24 / 10.17 | 0.01 / 0.01 | ~ 0 / ~ 0 |
| 24 | 3.80 / 3.90 | 6.42 | -0.76 / 1.02 | 5.43 / 13.68 | 0.06 / 0.06 | ~ 0 / ~ 0.21 |

### 3. Simulating behavioral responses

With the individual participant’s parameters at hand, we simulated behavioral responses for each participant in the following manner. For a given trial, an estimate of the probe stimulus orientation 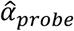 was drawn from a normal distribution centered on the nominal probe stimulus orientation of that trial with standard deviation *σ*_*S*_. Similarly, an estimate of the discrimination boundary orientation 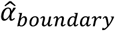 was drawn from a normal distribution centered on the nominal boundary orientation with standard deviation *σ*_*B*_. A response for the boundary discrimination judgment was computed by comparing 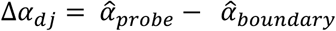 against criterion *τ*_*dj*_. For Δ*α*_*dj*_ > *τ*_*dj*_ the artificial observer responded “clockwise” otherwise it responded “counterclockwise”. However, with probability *λ*_*dj*_ the observer lapsed and gave a random response. Subsequently, an estimate of the reference stimulus orientation 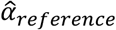 was drawn in a similar manner as for the probe stimulus. The response for the discrimination judgment was then computed by comparing 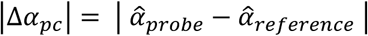 to criterion *τ*_*pc*_ of the perceptual comparison task. For |Δ*α*_*pc*_| ≤ *τ*_*pc*_ the observer responded “same”, and “different” otherwise. As for the discrimination judgement, the observer lapsed with probability *λ*_*pc*_, in which case it gave a random response. Low noise and high noise trials were simulated separately with their respective parameter estimates and the experiment was simulated 1000 times for each participant. Crucially, the artificial observer does not have an inbuilt reference repulsion bias, and any “error” responses that deviate from the nominally correct responses are due to lapses or noisy sampling of the true stimulus and boundary orientations.

### 4. Analyzing simulated data

The resulting simulated data was analyzed in the same way as the empirical data (see section *3.1.4.*). For each participant, *ΔPSEs* were taken as the trimmed mean (discarding 25% of the most extreme data points) of the distribution of the *ΔPSEs* simulated in 1000 iterations of the experiment.

### 5. Validating the artificial observer model

The artificial observer model described above underlies the assumption that the empirical data was generated by observers *without* a reference repulsion effect. In order for the comparison between simulated (baseline) biases and empirical biases to be valid, it is a is a crucial requirement that we could also accurately estimate the observer’s internal parameters, such as the variance in the stimulus sampling distributions, in the presence of a reference repulsion effect. If, however, the presence of a reference repulsion effect in the empirical data would bias the parameter estimation and would lead us to overestimate the contribution of trial-by-trial variability to the observed biases, a comparison between empirical and simulated (baseline) biases would not be valid and a conclusion about the presence or absence of reference repulsion could not be drawn. Therefore, we tested whether we could accurately separate biases introduced by reference repulsion on the one hand, and trial-by-trial fluctuations in the stimulus representations on the other hand, with our modeling approach. To this end, we first simulated an artificial observer with a reference repulsion effect. Subsequently, we estimated the observer’s parameters as described in section 2. Using the estimated parameters, we then simulated behavioral responses for an observer without a reference repulsion effect. As a result, we could compare the estimated parameters to the true generating parameters, and the simulated (baseline) bias of an observer without reference repulsion to the bias of the observer with reference repulsion. We simulated the observer with the generating parameters listed in **Table T2**. The observer was designed to have a small reference repulsion bias that deceased with increasing orientation difference between probe stimulus and discrimination boundary, in accordance with previous studies (Jazayeri & Movshon, 2007; Luu & Stocker, 2018; Experiment 1 of current study). In particular, the reference repulsion bias was computed on a trial-by-trial basis, based on the difference between the sampled boundary and probe stimulus orientation, 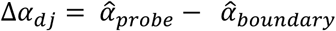. For low noise trials reference repulsion decreased linearly from 0.5 to 0 degrees over the range [0, 25] deg for Δ*α*_*dj*_, and was 0 for Δ*α*_*dj*_ > 25 *deg*. Analogously, for high noise trials it decreased from 2 to 0 degrees over the range [0, 25] deg for Δ*α*_*dj*_, and was 0 for Δ*α*_*dj*_ > 25 *deg*. We simulated the observer with reference repulsion once for each of the 18 stimulus sequences used in the experiment and estimated the parameters for each simulated dataset. For each instance, we then simulated the observer without reference repulsion 1000 times (see section 3). The simulated data for the observers with and without reference repulsion was analyzed in the same way as the empirical data (see also section 4).

**Table T2.**
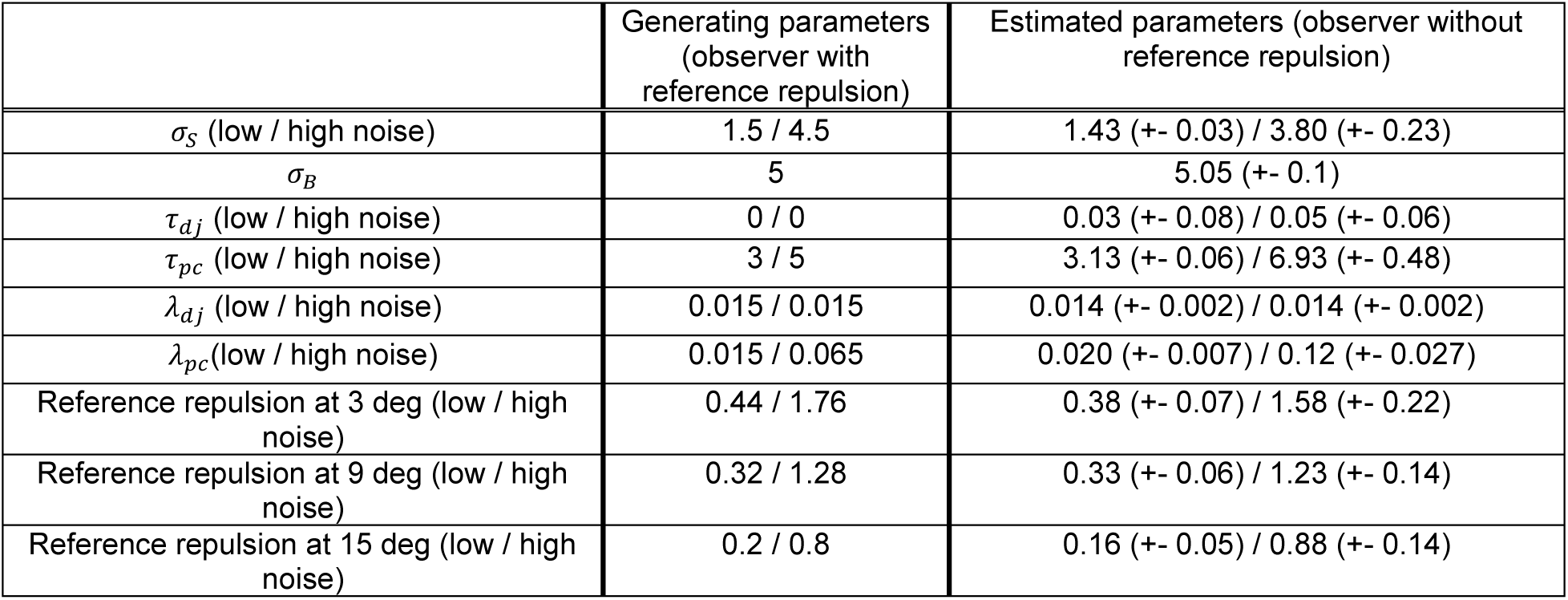
Parameters for observers with and without reference repulsion. Data was first simulated for an observer with reference repulsion, using the parameters in the first column. Subsequently, parameters were recovered from this data for an observer without reference repulsion. The estimated reference repulsion (right column) was computed as the difference between *ΔPSEs* for the observers with and without reference repulsion (**Figure S1**). Group means +-SEMs.

From the results shown in **Figure S1** and **Table T2**, one can see that a bias due to reference repulsion over and above a bias due to random trial-by-trial fluctuations in stimulus representations can be clearly detected. If anything, for an observer with reference repulsion the trial-by-trial variability in stimulus representations is slightly underestimated, and as a consequence the contribution of the reference repulsion effect to the overall bias is slightly overestimated. Since our simulation with exemplary parameters suggests that the analysis should be sensitive to even small reference repulsion biases, on the scale of 0.2 to 2 degrees, we are confident that it should be possible, in principle, to detect the presence of a reference repulsion bias.

## Variability in external stimuli versus internal stimulus representations

Our artificial observer model demonstrated that random fluctuations in external (physical) stimulus properties or internal stimulus representations could account for the apparent bias in our analysis for probe stimulus orientations close to the discrimination boundary in Experiment 2. Since we did not model external and internal variability separately, our model does not allow to estimate the individual contribution of these two sources of variability to the estimated bias. In order to assess whether the bias for probe stimuli oriented close to the discrimination boundary were mostly caused by internal or external variability we repeated our main analysis of Experiment 2 with the exception that the analysis was confined to trials in which: a) the participant made a correct boundary judgment, and b) *only including pairs of trials that had physically identical orientation stimuli*. Consequently, the difference in perceived orientation of the probe stimulus was computed between two identical sets of orientation stimuli, with the only difference between the sets being the orientation of the decision boundary and hence the direction of the boundary judgment. Therefore, any bias observed in this analysis should be either due to internal trial-by-trial fluctuations in stimulus representations or due to genuine perceptual biases. Consistent with our main analysis we found a highly significant bias away from the decision boundary (**Figure S2,** *F*(1,17) = 35.3, *p* = 1.6e-5). This bias was significantly stronger for the high noise compared to low noise stimuli (*F*(1,17) = 4.7, *p* = 0.044), but was not significantly different across boundary distances (*F*(1.24,21.08) = 0.08, *p* = 0.84, Greenhouse-Geisser corrected). Neither was there a significant interaction between noise and boundary distances (*F*(2,34) = 0.8, *p* = 0.47). Comparing magnitudes of biases found in this analysis to biases found in the main analysis (**Figure S2 solid vs dotted lines**), on average 99% of the bias for low noise stimuli and 89% of the bias for high noise stimuli can be attributed to internal fluctuations to physically matched stimuli. This indicates that most parts of the observed biases cannot be explained by trial-by-trial fluctuations in the orientation energy of the stimuli, but must be due to internal fluctuations in stimulus representations (bias close to discrimination boundary) or genuine repulsive perceptual biases (bias further away from discrimination boundary).

## Analyzing the bimodality of response distributions in Experiment 1

Since our Experiment 1 was similar to the “reference absent” condition of Experiment 1 by Zamboni et al. (2016), we were interested whether there would be evidence for reference repulsion in our Experiment 1, when analyzing the data similar to Zamboni et al. Strong reference repulsion should result in bimodal distributions of reproduction responses, since responses would be biased away from the discrimination boundary. Therefore, we quantified the deviation from unimodality of the response distributions with the descriptive dip-statistic (Hartigan & Hartigan, 1985). We computed the dip-statistic separately for low and high noise trials for each subject. Further, we combined data across all stimulus orientations by expressing the stimulus orientations relative to the discrimination boundary. A large dip-statistic provides evidence for deviations from a unimodal distribution (Hartigan & Hartigan, 1985). In order to statistically assess whether response distributions deviated from unimodality, we calculated 10,000 bootstrap estimates of the dip-statistic by sampling with replacement from the uniform distribution. A p-value was calculated by computing the proportion of bootstrap samples that yielded a larger dip-statistic than the one observed in the data. The alpha-threshold was set to 0.05. Similar to Zamboni et al., only one of twenty-four participants showed significantly multimodal response distributions in the current experiment (in both low and high noise conditions). Neither of the remaining 23 participants showed significantly multimodal distributions, neither in the low nor high noise condition. Similarly, the response histogram for the average observer does not show any sign of bimodality (**Figure S3**). This stands in contrast to our analyses of Experiment 1, which revealed small but significant reference repulsion biases. Together these findings suggest that the reference repulsion bias is present, but too small to lead to a clear bimodal pattern in the response distributions.

## Control Experiment – Order of boundary and perceptual comparison judgments

In Experiment 2, we instructed participants to make the boundary judgment (“Is the stimulus on the side of the discrimination boundary oriented more clockwise or counter-clockwise than the boundary?”) first and the perceptual comparison judgment (“Do the two orientation stimuli have the same or a different orientation?”) second. However, it is in principle possible that participants could have performed the two decisions in the opposite order. Since the order in which decisions were performed constitutes an important aspect of Experiment 2, we conducted a control experiment to investigate whether participants could have performed the decisions in the opposite order.

We previously hypothesized that making the boundary judgment first would constitute the more efficient order of decisions, since the boundary judgment required to use the previously memorized orientation of the discrimination boundary, which decays in memory and could be confounded/disrupted by processing the reference stimulus at the opposite side of fixation. Therefore, we predicted that participants would show worse performance on the task if they attempted to make the two decisions in the opposite order (same/different judgment first). In order to test this, we conducted a version of Experiment 2, in which on half of the blocks participants were instructed to perform the boundary judgment first (original instruction of Experiment 2), while on the other half of the blocks participants were instructed to perform the same/different judgment first. This allowed us to test whether our instructions about the order of the judgments were effective and to which extend participants could perform the task when making the same/different judgment first.

Seven naive participants (3 female/4 male, age range 19 - 30 years) took part in this control experiment. None of the participants took part in Experiments 1 to 3. All participants reported normal or corrected-to-normal vision and gave written, informed consent prior to the start of the study. The study was approved by the local ethical review board (CMO region Arnhem-Nijmegen, The Netherlands) and was in accordance with the Declaration of Helsinki.

The same experimental setup, stimulus parameters and procedures as in Experiment 2 were used, except that we only presented low noise stimuli and lowered the number of trials. Participants performed 12 blocks of 56 trials each, resulting in a total of 672 trials per participant, conducted in a single 1.5 to 2-hour session. Crucially, on half of the blocks participants were instructed to make the boundary judgment first, whereas on the other half they were instructed to make the same/different judgment first. The instructed order of judgments switched every 2 blocks and the starting order was counterbalanced across participants. In “boundary judgment first” blocks responses were given at the end of each trial by successively pressing the left/right arrow key for the boundary judgment and up/down arrow key for the same/different judgment. In “same/different judgment first” blocks the sequence of button presses was reversed. Similar to Experiment 2, at the beginning of the session, participants practiced the boundary judgment task in isolation in blocks of 36 trials until an acceptable performance level was reached. Before beginning the main experiment, participants practiced at least one “boundary judgment first” block and one “same/different judgment first” block of the main experiment, respectively.

We excluded one participant from the data analysis, since in the debriefing after the experiment she reported that she always performed the boundary judgment first, because she felt that she could not perform the boundary judgment after making the same/different judgment first. She reported that in the “same/different judgment first” blocks she merely reversed the order of button presses, but still performed the boundary judgment first.

The remaining participants showed a much higher accuracy in the boundary judgment task when instructed to make the boundary judgment first (75.55% +- 2.74 (SEM) correct boundary judgments) compared to when they were instructed to make this decision second (63.94% +- 3.04 (SEM) correct boundary judgments, see **Figure S4**). The difference in accuracy across conditions was statistically significant (*t*(5) = 4.9583, *p* = 0.0043). Moreover, the accuracy in the “boundary judgment first” blocks was similar to the accuracy for the low noise trials in Experiment 2 (Exp. 2: 77.12% correct boundary judgments).

In contrast, the same/different judgments appeared to be relatively unaffected by the order in which participants performed their judgments (see figure below). In particular, there was no significant difference in the width of the Gaussian models fit to the two conditions, indicating that participants’ representations underlying the same/different judgments were not strongly affected by the task order (“boundary judgment first”: 10.11°+- 1.32 (SEM); “same/different judgment first”: 9.31°+- 0.94 (SEM); “boundary judgment first” vs “same/different judgment first” *t*(5) = 1.9179, *p* = 0.1132).

These results indicate that the order in which participants make the decisions has important consequences for the performance in the boundary judgment task. Crucially, task performance is much higher when participants perform the boundary judgment first, i.e. the instructed order of Experiment 2. In contrast, when participants perform the same/different judgment first, their performance on the boundary task suffers substantially, suggesting that they could not maintain an accurate memory representation of the boundary while processing both orientation stimuli first. The substantially lowered accuracy compared to the “boundary judgment first” condition and compared to the performance in Experiment 2 suggests that participants did not perform the same/different judgment first in Experiment 2, but adhered to the instructed order of decisions.

---

**Figure S1:**
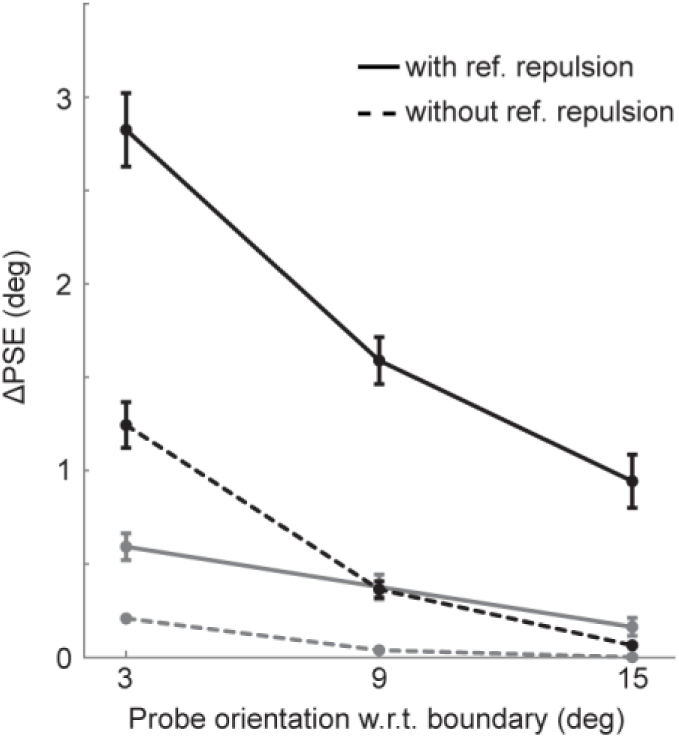
Simulated biases for artificial observers with (solid lines) and without (dotted lines) reference repulsion. The parameters underlying the observer without reference repulsion were estimated from the simulated data of the observer with reference repulsion (for parameters of both observers see **Table T2**). Biases are computed for low and high noise trials (black and grey data points). Biases of the observer with reference repulsion are clearly larger than biases of the observer without reference repulsion, indicating that reference repulsion effects could be detected if present in the data. Error bars represent SEMs over the simulations of the eighteen stimulus sequences used for the eighteen participants of Experiment 2. See also **Figure 4B**.

**Figure S2:**
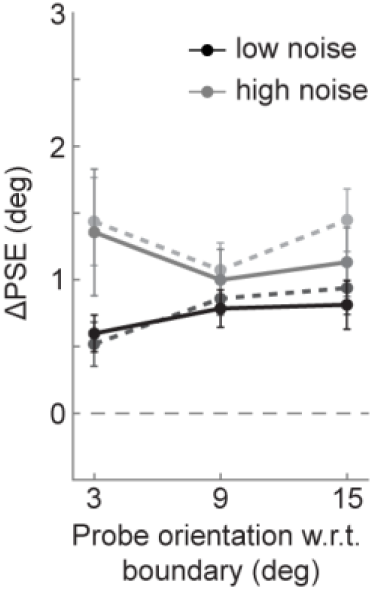
Biases computed for physically matched sets of orientation stimuli. In order to rule out that the observed biases revealed by the main analysis of Experiment 2 were caused by external fluctuations of the stochastically generated stimuli, we computed biases between pairs of trials with physically identical stimuli but opposite discrimination boundaries. Only trial pairs with correct responses were taken into account. Group bias for low and high noise trials (black and grey data points, solid lines) for all distances between probe and decision boundary (*x* axis) are shown. Most of the bias found in the main analysis (dotted lines, replotted from **Figure 4B)** persists when controlling for fluctuations in external, physical stimulus information. All data points represent group means and error bars depict SEMs.

**Figure S3:**
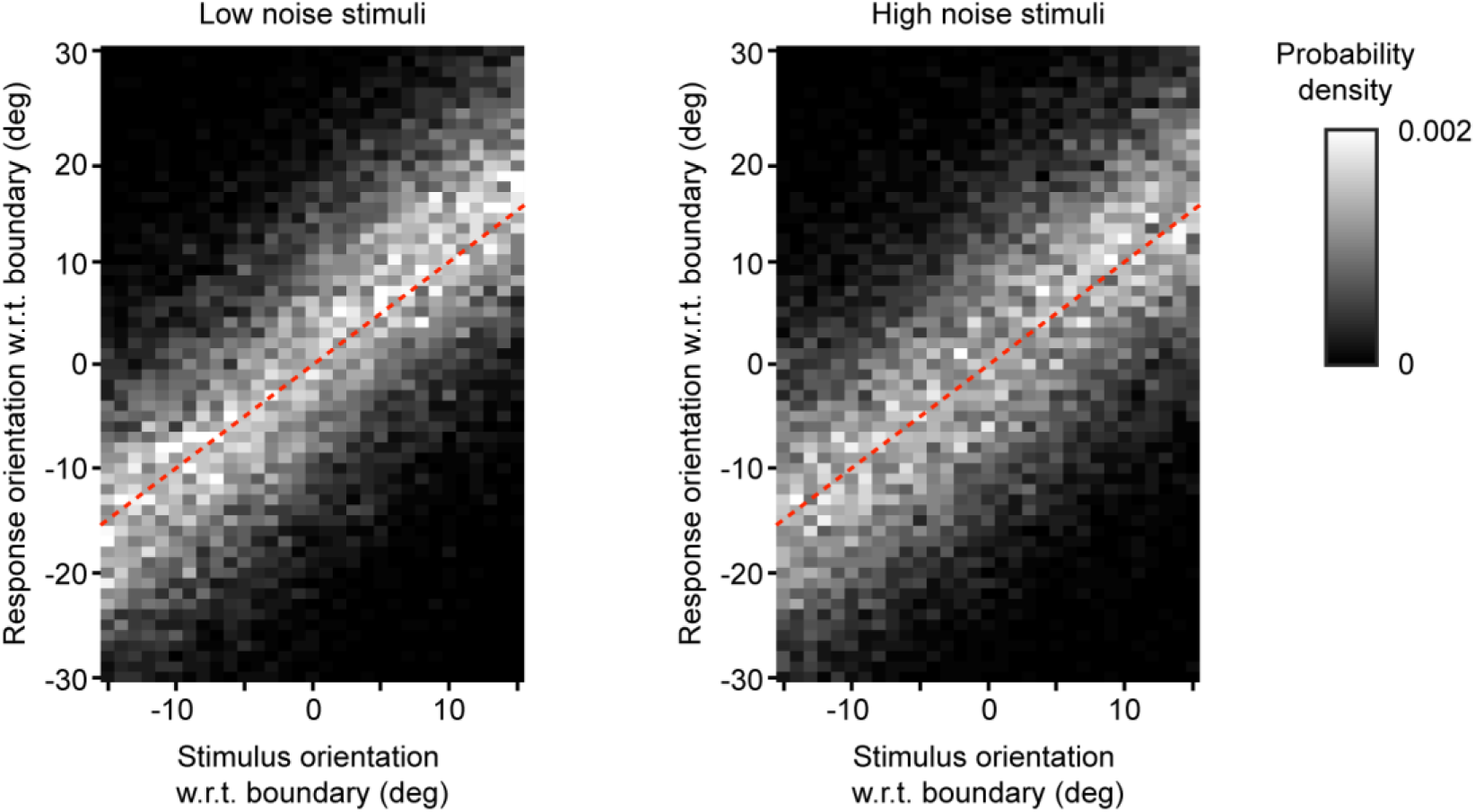
2-D histograms of reproduction responses of Experiment 1. The data were pooled across all observers and split in low (left panel) and high noise trials (right panel). Stimulus and response orientations were expressed relative to the orientation of the boundary (denoted by zero). Generally, reproductions responses matched the stimulus orientations well (red dotted lines indicate equal stimulus and response orientations). There was no indication for a bimodal distribution of reproduction responses, which would manifest as a horizontal band of decreased density around zero (compare to Figures 1 c-e in Jazayeri & Movshon (2007) and Figure 1d in Luu & Stocker (2018)). Importantly, despite the absence of a clear bimodal pattern of responses, we still found a highly significant repulsive bias in reproduction responses (see Figure 2B-D).

**Figure S4:**
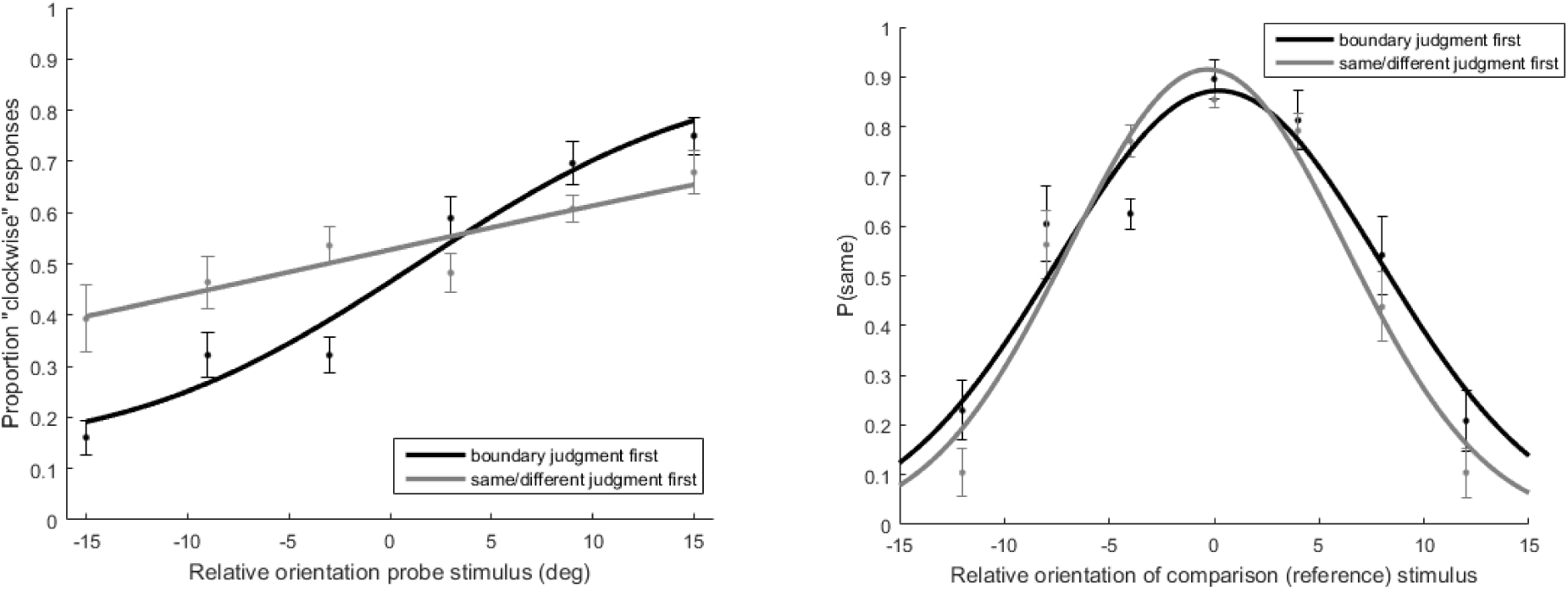
Left panel: Boundary judgment data of the average observer (n = 6) when instructed to make the boundary judgment first (black) or second (grey). Right panel: Response distribution for the perceptual comparison (same/different) judgment of the average observer, when making the boundary judgment first (black) or second (grey).

**Figure S5:**
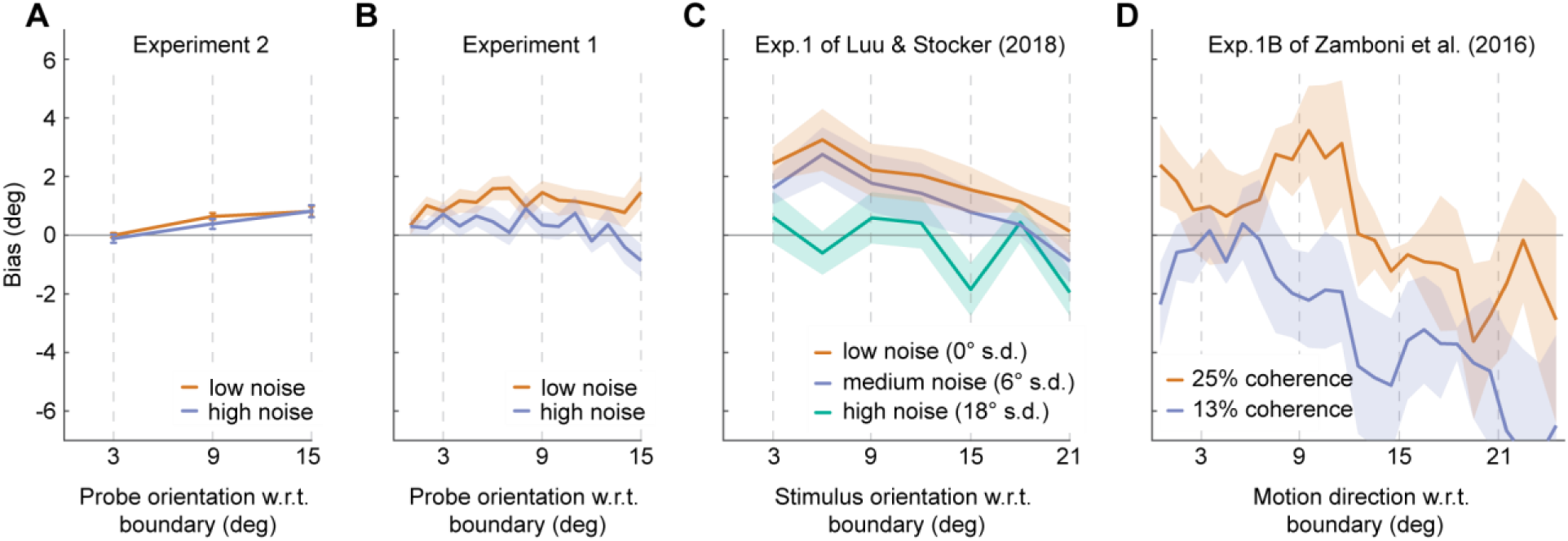
Comparison of data from four different experiments on reference repulsion. Biases are computed over all trials (i.e. including trials with correct and incorrect boundary judgments). **(A)** Results of Experiment 2 (reprint of **Figure 4C** in the main manuscript). This experiment measured the perceived orientation of a probe stimulus in direct comparison to a simultaneously presented reference stimulus. There was no perceptual bias for probe stimuli oriented close to the boundary, but a genuine repulsive perceptual bias for probe stimuli oriented further away from the boundary (for statistical tests see section 3.2 in the main manuscript). **(B)** Results of Experiment 1. This experiment measured the perceived orientation of the probe stimulus via orientation reproductions from memory and therefore is more susceptible to post-perceptual memory and decision biases. When statistically comparing Experiment 1 & 2, there were systematically larger repulsion biases in the reproduction responses in Experiment 1 when stimuli were oriented close (3°) to the boundary (two samples t-test for difference between Experiment 1 and 2; low noise: *t*(26.411) = 2.726, *p* = 0.011; high noise: *t*(29.869) = 2.142, *p* = 0.041). The difference in bias between Experiment 1 and 2 disappeared for stimulus orientations further away from the boundary (all *p >* 0.05), except for the high noise stimuli oriented 15° from the boundary for which the bias in Experiment 1 was significantly smaller than in Experiment 2 (*t*(28.827) = -2.791, *p* = 0.009). This analysis of all trials corroborates the analysis of only correct trials (see section 3.2 in the main manuscript), which indicates that post-perceptual reproduction responses of stimuli close to the discrimination boundary are systematically repelled from the boundary (Experiment 1), while the perceived orientation of those stimuli is unbiased (Experiment 2). However, the perceived orientations of probe stimuli oriented further away from the boundary are systematically biased away from the boundary. This stands in contrast to previous findings of reference repulsion, where the reference repulsion bias decreased with increasing distance from the boundary (see panels C and D). **(C)** Results of Experiment 1 by Luu & Stocker (2018) when computing biases over all (correct and incorrect) trials. The repulsion biases are maximal for orientations close to the boundary and decrease with increasing distance to the boundary (opposite to the pattern in panel A). **(D)** Results of Experiment 1B by Zamboni et al. (2016), in which the discrimination boundary was not presented during the reproduction phase. Similar to panel D, the repulsion bias decreases with increasing distance from the boundary. For noisier stimuli, the negative slope of the bias curve is preserved, but the repulsion bias for stimuli close to the boundary disappears in this analysis.

## References

Akaishi, R., Umeda, K., Nagase, A., & Sakai, K. (2014). Autonomous mechanism of internal choice estimate underlies decision inertia. Neuron, 81(1), 195–206, doi:10.1016/j.neuron.2013.10.018

Akrami, A., Kopec, C. D., Diamond, M. E., & Brody, C. D. (2018). Posterior parietal cortex represents sensory history and mediates its effects on behavior. Nature, 554, 368–372, doi:10.1038/nature25510

Bae, G.-Y., & Luck, S. J. (2017) Interactions between visual working memory representations. Attention, Perception, & Psychophysics, 79(8), 2376–2395, doi:10.3758/s13414-017-1404-8

Bliss, D. P., Sun, J. J., & D’Esposito M. (2017). Serial dependence is absent at the time of perception but increases in visual working memory. Scientific Reports, 7, Article number: 14739, doi:10.1038/s41598-017-15199-7

Brainard, D. H. (1997). The psychophysics toolbox. Spatial Vision, 10, 433–436.

Clifford, C. W. G. (2014). The tilt illusion: Phenomenology and functional implications. Vision Research, 104, 3–11, https://doi.org/10.1016/j.visres.2014.06.009

Corbett, J. E., Handy, T. C., & Enns, J. T. (2009). When do we know which way is up? The time course of orientation perception. Vision Research, 49, 28–37, https://doi.org/10.1016/j.visres.2008.09.020

Ding, S., Cueva, C. J., Tsodyks, M., & Qian, N. (2017). Visual perception as retrospective Bayesian decoding from high-to low-level features. Proceedings of the National Academy of Sciences, 114 (43) E9115–E9124, https://doi.org/10.1073/pnas.1706906114

Firestone, C., & Scholl, B. J. (2016). Cognition does not affect perception: evaluating the evidence for top-down effects. Behavioral and Brain Sciences, 39, e229, doi:10.1017/S0140525X15000965

Fritsche, M., Mostert, P., & de Lange, F. P. (2017). Opposite effects of recent history on perception and decision. Current Biology, 27(4), 590–595, doi:10.1016/j.cub.2017.01.006

Georgeson, M. A. (1973). Spatial Frequency Selectivity of a Visual Tilt Illusion. Nature, 245, 43–45, doi:10.1038/245043a0

Gibson, J. J. (1937). Adaptation, after-effect, and contrast in the perception of tilted lines. II. Simultaneous contrast and the areal restriction of the after-effect. Journal of Experimental Psychology, 20, 553–569.

Gibson, J. J., & Radner, M. (1937). Adaptation, after-effect, and contrast in the perception of tilted lines. I. Quantitative studies. Journal of Experimental Psychology 20, 453–467.

Huang, J., & Sekuler, R. (2010). Distortions in recall from visual memory: two classes of attractors at work. Journal of Vision, 10(2): 24, 1–27, http://journalofvision.org/10/2/24/, doi:10.1167/10.2.24.

Jazayeri, M., & Movshon, J. A. (2006). Optimal representation of sensory information by neural populations. Nature Neuroscience, 9(5), 690–696, doi:10.1038/nn1691

Jazayeri, M., & Movshon, J. A. (2007). A new perceptual illusion reveals mechanisms of sensory decoding. Nature, 446, 912–915, doi:10.1038/nature05739

Luu, L., & Stocker, A. A. (2018). Post-decision biases reveal a self-consistency principle in perceptual inference. eLife 7:e33334 doi: 10.7554/eLife.33334

Mohr, H. M., Linder, N. S., Dennis, H., & Sireteanu, R. (2011). Orientation-Specific Aftereffects to Mentally Generated Lines. Perception, 40(3), 272–290, https://doi.org/10.1068/p6781

Mulder, M. J., Wagenmakers, E.-J., Ratcliff, R., Boekel, W., & Forstmann, B. U. (2012). Bias in the brain: a diffusion model analysis of prior probability and potential payoff. Journal of Neuroscience, 32(7), 2335–2343, doi:10.1523/JNEUROSCI.4156-11.2012

Papadimitriou, C., Ferdoash, A., & Snyder, L. H. (2015) Ghosts in the machine: memory interference from the previous trial. Journal of Neurophysiology, 113(2), 567–577. http://doi.org/10.1152/jn.00402.2014

Pape, A.-A., & Siegel, M. (2016). Motor cortex activity predicts response alternation during sensorimotor decisions. Nature Communications, 7, 13098, doi:10.1038/ncomms13098

Prins, N., & Kingdom, F. A. A. (2009) Palamedes: Matlab routines for analyzing psychophysical data. http://www.palamedestoolbox.org

Rauber, H.-J., & Treue, S. (1998). Reference repulsion when judging the direction of visual motion. Perception, 27(4), 393–402, doi:10.1068/p270393

Saad, E., & Silvanto, J. (2013). How Visual Short-Term Memory Maintenance Modulates Subsequent Visual Aftereffects. Psychological Science, 24(5), 803–808, https://doi.org/10.1177/0956797612462140

Schneider, K. A., & Komlos, M. (2008). Attention biases decisions but does not alter appearance. Journal of Vision, 8(15):3. doi:10.1167/8.15.3.

Schwartz, O., Hsu, A., & Dayan P. (2007). Space and time in visual context. Nature Reviews Neuroscience, 8(7), 522–535, doi:10.1038/nrn2155

Scocchia, L., Cicchini, G. M., & Triesch, J. (2013). What’s “up”? Working memory contents can bias orientation processing, 78, 46–55, https://doi.org/10.1016/j.visres.2012.12.003

Snyder, J. S., Schwiedrzik, C. M., Vitela, A. D., & Melloni, L. (2015). How previous experience shapes perception in different sensory modalities. Frontiers in Human Neuroscience, 9, 594, doi:10.3389/fnhum.2015.00594

Stocker, A., & Simoncelli, E. P. (2008). A Bayesian model of conditioned perception. In Advances in Neural Information Processing Systems 20 (eds JC Platt, D Koller, Y Singer, ST Roweis), pp. 1409–1416. Curran Associates, Inc. (See http://papers.nips.cc/paper/3369-a-bayesian-model-of-conditionedperception.pdf.)

Summerfield, C., & de Lange, F. P. (2014). Expectation in perceptual decision making: neural and computational mechanisms. Nature Reviews Neuroscience, 15, 745–756, doi:10.1038/nrn3838

Tomassini, A., Morgan, M. J., & Solomon, J. A. (2010). Orientation uncertainty reduces perceived obliquity. Vision Research, 50(5), 541–547, https://doi.org/10.1016/j.visres.2009.12.005

Visscher, K. M., Kahana, M. J., & Sekuler, R. (2009). Trial-to-trial carryover in auditory short-term memory. Journal of Experimental Psychology: Learning, Memory, and Cognition, 35(1), 46–56, doi:10.1037/a0013412

Webster, M.A. (2015). Visual adaptation. Annual Review of Vision Science, 1, 547–567, doi:10.1146/annurev-vision-082114-035509

Wei, X.-X., & Stocker, A. A. (2015). A Bayesian observer model constrained by efficient coding can explain ‘anti-Bayesian’ percepts. Nature Neuroscience 18, 1509–1517, doi:10.1038/nn.4105

Wenderoth, P., & Johnstone, S. (1987). Possible neural substrates for orientation analysis and perception. Perception, 16, 693–709, https://doi.org/10.1068/p160693

Zamboni, E., Ledgeway, T., McGraw, P. V., & Schluppeck, D. (2016). Do perceptual biases emerge early or late in visual processing? Decision-biases in motion perception. Proceedings of the Royal Society B: Biological Sciences, 283(1833), 20160263, doi:10.1098/rspb.2016.0263

## Supplemental references

Hartigan JA, Hartigan PM. (1985). The dip test of unimodality. Ann. Stat.13. doi:10.1214/aos/1176346577

Zamboni E., Ledgeway T., McGraw P.V., & Schluppeck D (2016) Data from: Do perceptual biases emerge early or late in visual processing? Decision-biases in motion perception. Dryad Digital Repository. https://doi.org/10.5061/dryad.ms84h

